# Neural dynamics of induced throat vibrations during vocal emotion recognition

**DOI:** 10.1101/2025.03.24.644945

**Authors:** Damien Benis, Garance Selosse, Manuela Filippa, Jennifer Greilsamer, Ben Meuleman, Didier Grandjean, Leonardo Ceravolo

## Abstract

Despite a large literature on the psychological and brain mechanisms of emotional prosody perception, the perspective of embodied cognition on these mechanisms has been largely neglected. Here we investigated the influence of induced bodily vibrations near the vocal tract on the categorization of emotionally ambiguous vocalizations while acquiring event-related potentials (ERPs) through electroencephalography. Emotional voices were morphed between a fearful expression and the speaker’s identity-matching angry expression, creating blends of emotions in each voice. In line with our hypotheses, behavioral results revealed that induced vibrations biased the participants’ emotion categorizations. Namely, induced throat vibrations of anger shifted responses toward anger for the most ambiguous voices, while fear vibrations shifted responses toward fear for anger-leaning voices. ERPs indicated that both anger and fear vibrations modulated frontocentral amplitude from about 75 ms, with the sign of the effect depending on time bins and morphing levels. These modulations show that vibrations are registered from early stages of processing. Source reconstruction highlighted effects of vibration types in prefrontal, cingulate, parietal, and medial and lateral temporal cortices, and to a lesser extent in somatosensory and insular cortices. Our results suggest that voice-associated vibrations may contribute to vocal emotion processing through an embodied feedback mechanism.

## 1 Introduction

Verbal and non-verbal communication is essential for humans, and voice tone or “emotional prosody” represents the temporal fluctuations of the pitch, amplitude and spectrum (Grandjean et al., 2006; Laukka et al., 2005; Risberg & Lubker, 1978; Scherer & Oshinsky, 1977). Processing these variations allows listeners to subjectively attribute the emotional state of speakers, influenced by context (Brunswik, 1956; Grandjean et al., 2006; Scherer, 2003). One key mechanism thought to underlie this attribution is the embodied ‘simulation’ of what we hear (Hawk et al., 2012). Also named “emotional embodiment”, it is defined as the motor, sensory and perceptual reexperiencing or simulating of the emotion perceived in a peer— framework of the Simulation of Smiles (SIMS) model (Niedenthal, 2007; Niedenthal et al., 2010). This process has primarily been described in the visual modality, with the perception of facial expressions in others generating facial mimicry in the observer—stronger for emotional vs. non-emotional faces (Chartrand & Bargh, 1999; Dimberg, 1982; Korb et al., 2010; Künecke et al., 2017; Moody & McIntosh, 2011; Moody et al., 2007). Recent studies have supported the idea that facial mimicry represents context-dependent sensorimotor simulation of an observed emotion rather than a mere muscular reproduction of an observed facial expression (Borgomaneri et al., 2020; Drimalla et al., 2019; Hess & Blairy, 2001; Hess & Fischer, 2014; Philip et al., 2018). Importantly, some studies suggest that this process may impact the accurate decoding of facial emotional information especially in women (Hyniewska & Sato, 2015; Korb et al., 2010; Schneider et al., 2013; Stel & van Knippenberg, 2008), although not exclusively (Blairy et al., 1999; Hess & Blairy, 2001; Holland et al., 2021).

Studies on the contribution of embodiment in the field of emotional prosody decoding are scarcer than in the visual domain. Studies showed that passive listening of speech signals elicits activations in motor regions of the brain similar to those involved during speech production (Fadiga et al., 2002; Watkins et al., 2003; Wilson et al., 2004). Facilitated language comprehension when the primary motor areas controlling the lips/tongue were stimulated transcranially was also observed (D’Ausilio et al., 2009). One mechanism underlying emotion embodiment in voice signals may be body resonances or ‘vibrations’. In fact, vibrations—especially close to the vocal folds—may represent a reliable interoceptive and/or exteroceptive feedback during vocal emotion expression (Filippa et al., 2026). Indeed, during voice expression, vibrations from the vocal cords are transmitted through the speaker’s skin and body tissues (Munger & Thomson, 2008; Švec et al., 2005), with a frequency corresponding to the fundamental frequency of the produced auditory tone (Sundberg, 1992). These bodily vibrations would constitute, for the speaker, a dynamic feedback on several characteristics of his/her own voice (Sundberg, 1992). Lylykangas and colleagues (2009) highlighted the potential to successfully use vibrotactile feedback as relevant information for speed regulation messages. Similarly, Tuuri and colleagues (2010) compared audio and tactile feedback from the same speech stimuli as speed regulation cues and found that the information was delivered in a similar manner across modalities. However, there is still a paucity of research in this field and we intend to fill this gap. To investigate precise dynamic neural correlates of the embodied account of vocal emotion perception concurrent to induced throat vibrations, we used surface electroencephalography (EEG) using event-related potentials (ERPs) as the underlying dependent measure.

ERP correlates of emotional prosody decoding have been extensively studied, with evidence for a three-stage processing (Schirmer & Kotz, 2006). In line with this model, a wealth of ERP studies has shown the implication of several early and late ERP components in the processing of emotional vocalizations. Early components underlie auditory-sensory processing, attention and early recognition of meaning around 100ms after stimulus onset (N100, frontocentral area). It might reflect early low-level prosody analysis and emotion discrimination (Gädeke et al., 2013; Paquette et al., 2020; Tarai & Srinivasan, 2019). Basic emotions are also distinguished in the frontal/parietal P200 component (Duville et al., 2022; Gädeke et al., 2013; Maltezou-Papastylianou et al., 2022; Paulmann et al., 2013; Paulmann & Uskul, 2017). The P200 is a positive deflection around 200 milliseconds reflecting increased attention to emotional stimuli for facilitated processing (Paulmann & Kotz, 2008; Thomas et al., 2007). Later components are commonly associated with higher-level mechanisms of emotional prosody processing—appraisal, contextual relevance and mental representations (Duville et al., 2022; Pell & Kotz, 2021). This includes ∼300-400 milliseconds deflections (P300/N400) in frontal/frontocentral areas during emotional prosody processing (Carminati et al., 2018; Proverbio et al., 2020; Tarai & Srinivasan, 2019), underlying deviant auditory stimuli detection (Lindín et al., 2004; Tsolaki et al., 2015). Finally, the Late Positive Component (LPC) is a positive deflection observed at a later stage in frontal/frontocentral and parietal areas, associated with higher-order cognitive processes such as arousal information (Paulmann et al., 2013), expectancy violation in prosody (Astésano et al., 2004; Brattico et al., 2006; Paulmann et al., 2012) and emotional ‘meaning’ (Wei et al., 2022; Zora et al., 2020). Mismatching prosody stimuli also elicits an LPC, potentially indicating an integration process of the different sources of information (Brattico et al., 2006; Paulmann et al., 2012).

Concerning functional MRI studies, processing emotional prosody activates subcortical and cortical networks (Ceravolo et al., 2021; Grandjean, 2021; Peron et al., 2016; Witteman et al., 2012), including the bilateral superior temporal, inferior frontal (Ceravolo et al., 2026) and orbitofrontal cortex, cerebellum, etc. Additionally, the motor cortex is also involved during speech perception (Fadiga et al., 2002; Watkins et al., 2003) while interoceptive signals trigger insular activity, a region responsible for the integration of internal and external physiological signals and emotional processes (Adolfi et al., 2017; Casals-Gutierrez & Abbey, 2020). The somatosensory cortex or the supplementary motor area (SMA) also play a critical role in interoceptive and proprioceptive processing (Engelen et al., 2023; Gibson, 2019; Khalsa et al., 2018).

Our study aims to explore the potential interoceptive and/or exteroceptive feedbacks represented by bodily resonances—induced external throat vibrations, and their impact on emotional prosody recognition. Revealing such a link between vocal emotion perception and induced vibrations would constitute convincing, causal evidence for a crucial role of vibrotactile feedback as a physiological mechanism of vocal emotion embodiment. We recently demonstrated the neural correlates of measured throat vibrations during emotional voice expression (Selosse et al., 2025) and our results are coherent with above mentioned literature. They indeed highlight that voice-generated throat vibrations recruit brain regions associated both with the expression and perception of vocal emotion, and interoception. Associations between interoception/proprioception and emotional processing were previously studied but never using vibrations concurrent to expressed voice signals (Burleson & Quigley, 2021; Couto et al., 2015; Critchley & Garfinkel, 2017). Other related work includes: interoceptive accuracy (Mendoza-Medialdea & Ruiz-Padial, 2021), ERPs of interoception in emotion processing (P300 component; Herbert et al., 2007; Pollatos et al., 2005), a somatosensory-specific P300 component (parietal regions) involved in somatosensory perception (Al et al., 2020; Auksztulewicz & Blankenburg, 2013; Has et al., 2021), interoceptive priming on faces (Salamone et al., 2021). In addition, other studies further support the idea that emotionally relevant stimuli can modulate attention and early perceptual processing across modalities (Brosch, Grandjean, et al., 2008; Brosch, Sander, et al., 2008)— in the case of the present study, auditory and vibrotactile modalities.

This design directly builds on an earlier behavioral test of the same mechanism from our group (Selosse et al., 2022), which used the same vibrator and throat placement to test whether congruent vibrations would facilitate the categorization of joy and anger vocalizations varying in acoustic intensity. That study found that vibrations increased confusion and decreased discrimination and accuracy, with no significant interaction between vibration and stimulus ambiguity—a result at odds with the embodied-feedback hypothesis as originally framed. Critically, that paper’s own discussion noted that intensity-based ambiguity may not adequately capture perceptual ambiguity, and proposed that morphing two distinct emotions into graded blends would constitute a more suitable test. The present study implements this design with a forced binary categorization task, allowing us to isolate a directional bias from the general perceptual disruption reported previously.

Therefore, and to the best of our knowledge, no study has specifically investigated the contribution of throat vibrations to the auditory perception of emotions using EEG. The present study aims to further explore this mechanism. The main mechanistic rationale is that vocal cords vibrations, seen as a ‘mechanical’ consequence of the production of vocalized signals, would actually represent a feedback that could impact the auditory perception and recognition of vocal emotions. Namely, according to an embodied perception perspective: as a valid feedback, vibratory signals would influence voice production for the speaker and should therefore also have an impact on perception if induced in the listener. At the behavioral level, we first postulated that: a) vibrations would affect the discriminative perceptual processes of emotional vocalizations (more frequent categorization of the morphed voice as “anger” when the vibrations also transmit anger and likewise for fear) or when compared to the ‘no-vibration’ condition; b) these effects would occur especially for the most ambiguous stimuli (i.e., voices containing 40 to 60% of each emotion), as these would trigger sensory information seeking for clarification (Niedenthal, 2007; Niedenthal et al., 2010), here induced throat vibrations. At the neural level, we postulated: c) vibration-specific ERPs modulations in early-to-intermediate timebins, modulated by interoceptive/proprioceptive processes (Herbert et al., 2007; Mendoza-Medialdea & Ruiz-Padial, 2021; Pollatos et al., 2005) with source modulations in the insular, somatosensory, prefrontal, and cingulate cortices as well as in the SMA/pre-SMA (Adolfi et al., 2017; Casals-Gutierrez & Abbey, 2020; Engelen et al., 2023; Gibson, 2019; Khalsa et al., 2018), for anger or fear vibrations compared to no-vibration.

## 2 Method

### 2.1 Participants

Sample size was decided a priori based on repeated measures MANOVA in ‘G*Power 3.1’ software, with: effect size=.25, power=.80, alpha=.05, number of measures=15 and correlation among repeated measures=0.5 (Faul et al., 2007). This approximation—the closest possible to mixed-effects modeling—resulted in a required sample size of N=24. We therefore recruited twenty-four healthy participants (18 females and 6 males; Age=18-41; *M_Age_*=23.67; *SD_Age_*=5.11; Participant gender was entered as a fixed effect, see below). Data from all participants were used in the analyses as no exclusions were necessary. Participants had normal hearing, no neurologic or psychiatric history and gave informed written consent. The study was approved by the state-wise (Geneva canton) ethics committee (CCER nr. 2020-02262) and conducted according to the declaration of Helsinki.

### 2.2 Material

Voices (‘Aah’, Montreal affective voices database; Belin et al., 2008) were presented through headphones (HD25, Sennheiser, DE) and morphed on the emotion dimension (anger, fear) using Matlab (The Mathworks Inc., Natick, MA, USA) and the PsychToolbox (Brainard, 1997; Kleiner et al., 2007; Pelli, 1997). Morphing fearful expression with the speaker’s identity-matching angry expression resulted in five conditions of 10 voices each [10% fear-90% anger (FA_10/90_), 40% fear-60% anger (FA_40/60_), 50% fear-50% anger (FA_50/50_), 60% fear-40% anger (FA_60/40_), and 90% fear-10% anger (FA_90/10_)], using the STRAIGHT toolbox (Kawahara & Matsui, 2003). See Figure 1**C**. Voice stimuli and vibrations were acoustically assessed (amplitude, pitch, loudness, duration, spectral centroid and slope, F1 frequency; Supplementary Table 1 & 2, respectively). Morphing conditions and actors were randomly distributed across trials. The choice of using anger and fear was due to the fact that these emotions are among the most biologically-relevant for survival.

**Figure 1.**
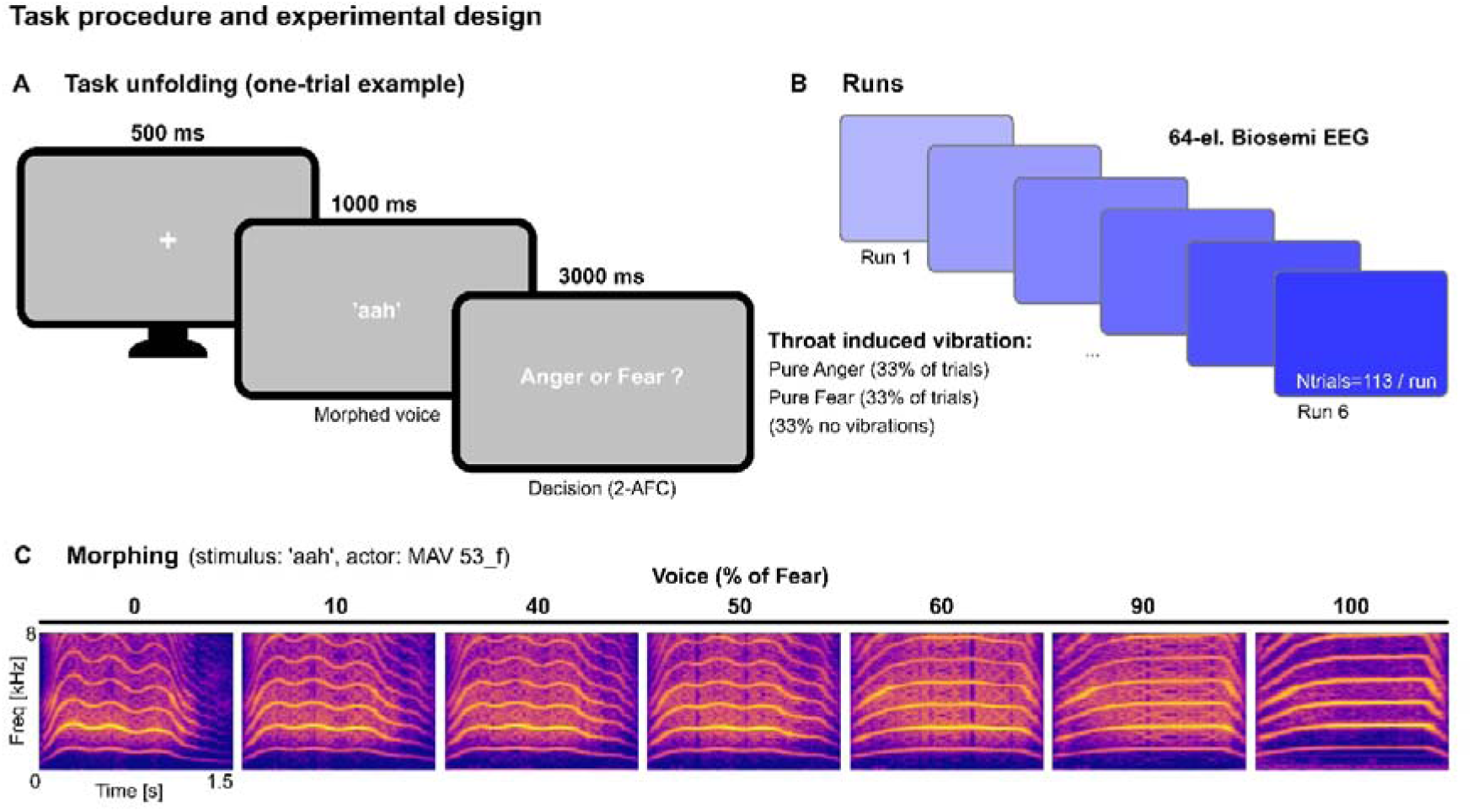
Experimental design and stimuli. (**A**) Participants were instructed to focus on a central fixation cross displayed on a grey screen for five hundred milliseconds. Emotional voice stimuli were presented through headphones for one second. Vibrations were induced near the vocal tract, on the skin of the throat at the same time in two thirds of the trials and expressed pure anger or pure fear. After each voice stimulus, the participant had to categorize the emotion expressed auditorily using the response keys, as fast and as accurately as possible and within the 3-seconds response window. (**B**) The task was split into 6 runs of 113 trials each, for a grand total of 678 trials per participant, while surface EEG was acquired through a 64-electrode Biosemi system. (**C**) Stimuli from the Montreal Affective Voices database were used, and morphed with actor-matching on an anger-fear continuum in steps of 10%. For the sake of task duration, morphing levels retained included anger-fear and fear-anger stimuli morphed 10%, 40%, 50%, 60%, and 90%.

Throat vibrations were induced using a small vibrator (BC-10, Ortofon, DK; dimensions: 13.5*29.5*18.0mm, weight: 16.5gr, sensitivity: 118dB, harmonic distortion: 1.5%, impedance: 15Ω, frequency range: 100-1000Hz, sampling rate: 1000Hz). These expressed pure anger or pure fear in 2/3^rd^ of the trials (unmorphed ‘Aah’; anger, fear) while no vibrations were induced in the remaining trials (control, 1/3^rd^). Participants confirmed upon debriefing that vibrations were not audible, even without the headphones. The vibrator was placed close to the vocal cords, taped on the left side to the laryngeal prominence with kinesiology tape—soft and able to stretch while being strong and highly adhesive. This location fits our general hypothesis that throat vibrations may be used as a vibrotactile feedback. It also matches relevant locations in this context (Munger & Thomson, 2008; Nolan et al., 2009; Sundberg, 1992). Whenever a vibration was induced, audio and vibrator actor were the same identity for the sake of consistency and to avoid intonation—and interpretation—issues. There were 20 stimuli for the vibrations (two emotions, ten actors).

### 2.3 Task Procedure

The experiment comprised six runs (three fear-induced, three anger-induced vibrations; no-vibration trials were present in each run, representing 33% of the total trials count; run order counterbalanced) of N=113 randomly presented trials each (Figure 1**B**), for a grand total of 678 trials. Each trial started with a fixation cross (500ms), followed by a voice stimulus (1000ms), then explicit emotion categorization for 3000ms (buttons: anger, fear; key labels counterbalanced between participants; Figure 1**A**). Participants were instructed to answer as fast and as accurately as possible. Vibrations were induced simultaneously to the voices. The final factorial design is a 5 Morphing (FA_10/90_, FA_40/60_, FA_50/50_, FA_60/40_, FA_90/10_) by 3 Vibration (No Vibration, Anger, Fear) within-participant design.

### 2.4 Statistical Analysis

#### 2.4.1 Trial-level behavioral analysis

Responses were coded anger=0, fear=1 and analyzed with a binomial generalized linear mixed model computed in R 4.2.0 through Rstudio (Venables & Smith, 2003) to analyze the responses as a function of fixed effects Vibration (anger, fear, none) and Morphing (10, 40, 50, 60, 90% fear morphing), their interaction, participant gender and speaker gender. Random effects were uncorrelated by-participant slopes for Vibration and Morphing, and random intercepts for run and stimulus identity (see formula below).

Reaction times were analyzed with the same structure in a Gaussian linear mixed model (lmer); they showed no significant effects and are reported in Supplementary Table 3. Both models were computed using the ‘lme4’ package (Bates et al., 2015) using Wald χ² tests (car::Anova, type II).

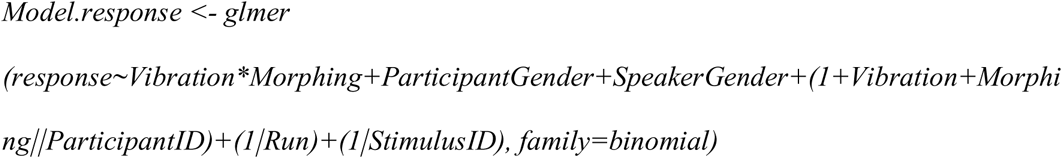

Planned contrasts were estimated marginal means (emmeans; Lenth, 2021) on the log-odds scale and are reported as odds ratios with 95% confidence intervals, which serve as the effect sizes. For each morphing level we compared the two vibration pairs (anger vs none, fear vs none), with Dunnett correction (family of two comparison) within level.

Analyzed data excluded trials where no response was given, namely 96 trials across participants on a total of 16,272 trials, representing only 0.59% of exclusions.

#### 2.4.2 EEG preprocessing

Surface EEG was acquired with 64 amplified electrodes (Biosemi ActiveThree system, BioSemi B.V., Netherlands) at a sampling rate of 2048Hz, preprocessing was performed using Brainvision analyser 2.2 (Brain Products, Munich, Germany) and Fieldtrip (Oostenveld et al., 2011). Spline interpolation was applied to artifacted channels (5.34±4.74% of channels) and an average reference was applied. Blink and saccade artifacts were rejected using independent component analysis. Raw data were split into two-seconds epochs (-1s to +1s) centered on voice onset. Baseline window was 1000 ms before voice onset. Peak-to-peak amplitude was calculated for all epochs across all channels, and a trial rejection threshold was calculated as the median +1.96 standard deviation of the obtained peak-to-peak amplitude distribution (∼114μv). Rejection was applied channel by channel, so the number of retained trials can differ between channels of the same participant. For display, ERPs were averaged per participant, channel, Morphing and Vibration, then across participants (mean trials per participant and condition: 40.95 ± 6.08; Supplementary Table 7). Statistics used single trials (2.4.3). Topographical clustering methods are reported in the supplementary material (‘Supplementary EEG methods’, *I.*).

#### 2.4.3 ERP statistical analysis

Statistics were run on single-trial amplitude (µV, baseline-corrected) averaged within 20 consecutive 50-ms bins from 0 to 1000 ms after voice onset (bin centres 25-975 ms), for trials with a response (99.41% of all trials). Four scalp clusters (C1 posterior, C2 centro-temporal, C3 anterior-lateral frontal, C4 frontocentral; electrode lists in Supplementary Figure 2) defined the regions of interest. For each cluster and each time bin separately, we fitted a Bayesian Gaussian linear mixed model (brms; Bürkner, 2017) with Vibration * Morphing, participant gender and speaker gender as fixed effects, and random intercepts for participant and EEG channel. Factors were sum-coded. A trial-level random intercept was not included because the models did not converge reliably with it. Trial-level data across electrodes made frequentist tests significant in all 20 bins in every cluster, so we used Bayes factors. (Supplementary EEG methods II.)

We used weakly informative priors on the µV scale: normal(0, 10) for fixed effects and Student-t(3, 0, 10) for the standard deviations of random effects and residuals. Each model ran four chains of 2000 iterations (500 warm-up; seed 1). Convergence was satisfactory in all 80 full models (4 clusters * 20 bins; R-hat ≤ 1.012; bulk ESS ≥ 610, tail ESS ≥ 1016).

Evidence for the Vibration * Morphing interaction was quantified with a Bayes factor (BF) comparing the full model with the same model without the interaction, estimated by bridge sampling (bridgesampling package). We labelled BF (K) following Jeffreys and Raftery (Jeffreys, 1998; Kass & Raftery, 1995): 3.2 < BF < 10 (+), 10 < BF < 100 (++), BF > 100 (+++); BF below 3.2 was not treated as evidence for an interaction. In bins with BF ≥ 3.2 we computed estimated marginal means from the posterior draws (emmeans) and the two vibration contrasts within each morphing level (anger vs. none, fear vs. none), reported as posterior median and 95% highest-posterior-density (HPD) interval. A contrast is called ‘credibly non-zero’ when its 95% HPD interval excludes zero. No further correction for multiple comparisons was applied across bins, clusters or contrasts; contrasts are interpreted only in bins where the interaction BF supports an effect.

#### 2.4.4 Source reconstruction

Cortical sources were estimated with a linear-constrained minimum-variance (LCMV) beamformer on the preprocessed epochs. Source activity was summarised in the 360 cortical parcels of the HCP-MMP1 atlas (Glasser et al., 2016) and restricted to the eight 50-ms windows with a supported interaction in C4 (75, 325, 475, 525, 625, 775, 825 and 925 ms). We used a sequential analysis approach to detect significant vibration effects for each source label in each time-window with a supported interaction in C4.

For each parcel and window, amplitude was analysed with a Gaussian mixed model with Morphing × Vibration as fixed effects and a random intercept for participant. Values beyond three interquartile ranges were excluded as outliers.

If, in a parcel and time window, Morphing × Vibration interaction reached significance, joint omnibus test were performed testing the effect of vibration for each morphing level (Benjamini–Hochberg method FDR correction for each omnibus test, 173 significant ROIs/Timewindows X 5 morphing levels = 865 tests). If joint-omnibus tests reached significance, contrasts were computed without emmeans adjustment, and the *p*-values of all anger-vs-none and fear-vs-none contrasts were adjusted together with the Benjamini– Hochberg method (false discovery rate) Estimated marginal means (emmeans) gave the contrasts anger − none and fear − none within each morphing level, expressed as z ratios (positive: stronger activity with vibration than without). P-values were adjusted by the Benjamini–Hochberg method (false discovery rate, FDR) using adjusted *p*FDR<.05. Because windows were selected from the ERPs, these tests describe where the vibration effects localize and are not independent confirmations. Figure 4 shows the surviving contrasts and the colorbar saturates at *z*=±[4.5]. See Supplementary EEG methods III.

## 3 Results

In this study, the participants had to categorize morphed voice emotions concomitant to induced throat vibrations in 2/3^rd^ of the trials. We analyzed the influence of these vibrations on voice recognition responses, ERPs and source modulations in the brain.

### 3.1 Behavioral responses

According to our research question, we first tested whether external vocal tract induced vibrations would influence the categorization of perceived voices. Namely, that vibrations inducing anger would shift the response curve and that incidentally voices, especially the most ambiguous ones, would be evaluated as expressing more anger. Similar congruent results were expected for fear.

Both induced vibrations (χ*^2^*(2)=8.27, *p*=.016) and voice emotion (χ*^2^*(4)=17.76, *p*=.001; Supplementary Figure 1) influenced categorization and their interaction was also significant (χ*^2^*(8)=23.96, *p*=.002). Model effect size was small to medium (Fixed effects R^2^=.14; Fixed and random effects R^2^=.36). See Figure 2 for illustration and Supplementary Table 3 for all effects. The modest voice effect reflects the inclusion of stimulus identity as a random effect.

**Figure 2.**
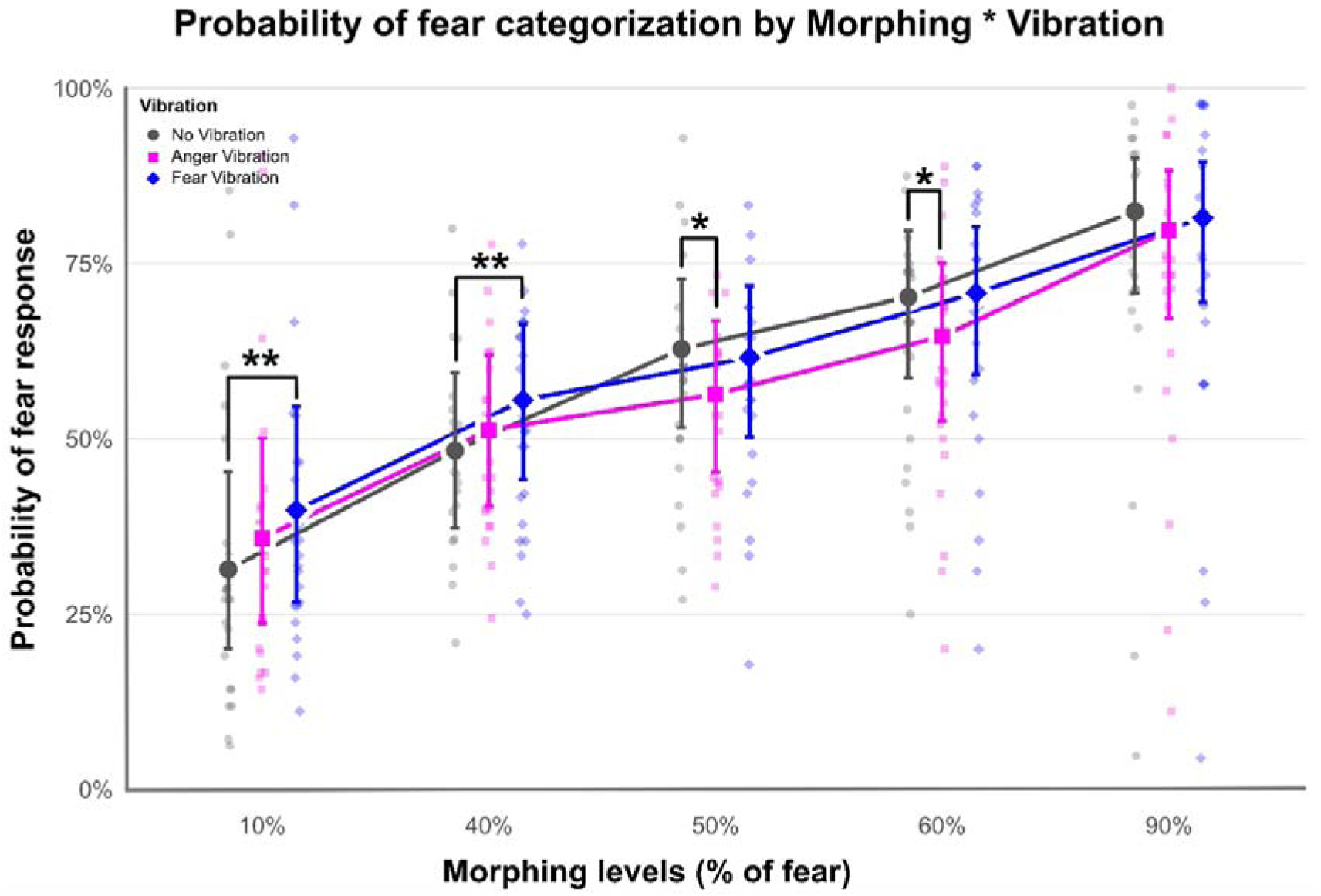
Effect of Vibration on the probability of fear responses, per morphing level. The probability of a ‘fear’ response—the inverse pattern of an ‘anger’ response—is illustrated on the Y axis per morphing levels (X axis). Vibration conditions are colored: grey, no vibration (33% of trials); pink, anger vibrations (33% of trials); blue, fear vibrations (33% of trials). Points represent individual values per Morphing and Vibration levels, error bars represent the 95% confidence interval. * *p*<.05, ** *p*<.01.

To test the effect of vibrations at each morphing level, we computed planned comparisons based on the interaction (Dunnett-adjusted; odds ratios, OR, refer to the odds of a ‘fear’ response, vibration relative to no vibration; Figure 2). Fear vibration increased fear responses for the least ambiguous voice, with 10% fear (90% anger; OR=1.449, SE=0.1600, *z*=3.359, *p*=.0015), and for the ambiguous voice with 40% fear (OR=1.336, SE=0.1300, *z*=2.972, *p*=.0058). Anger vibration, by contrast, shifted responses toward anger only within the ambiguous range, at 50% fear (OR=0.764, SE=0.0786, *z*=-2.617, *p*=.0172) and 60% fear (OR=0.772, SE=0.0818, *z*=-2.443, *p*=.0280). Neither vibration changed responses to the most fearful voice (90% fear; all *p*≥.26). Fear vibration therefore influenced categorization from the least ambiguous voice into the ambiguous range, whereas anger vibration influenced the most ambiguous voices only: as predicted for anger, and beyond our prediction for fear. Exhaustive post hoc comparisons are reported in Supplementary Table 8.

### 3.2 EEG results

#### 3.2.1 Event-Related Potentials

Bayesian mixed models tested the Vibration * Morphing interaction separately for each topographical cluster and 50-ms timebin (Section 2.4.3). In the frontocentral cluster C4 (Supplementary Figure 2), the interaction was supported (Bayes factor *K*>3.2) in 8 of the 20 timebins, with decisive evidence (*K*>100) in 7 of them (all except 525 ms). The other clusters showed far less evidence: one timebin in C3 (75 ms, 3.2<*K*<10), none in C2, and three in C1 (75, 825 and 925 ms, all *K*>100, the same bins as in C4). Because C4 showed evidence in the most timebins, we describe it here; the other clusters are reported in the Supplementary Tables 4 and 6 and Supplementary EEG results I and II.

The C4 grand-average ERP showed a small positive deflection at 100-150 ms (P100), a large positive deflection peaking at about 275 ms (P200) and a sustained, negative deflection from about 500 ms after stimulus onset (Figure 3**A**). Evidence for the interaction was decisive at 50-100, 300-350, 450-500, 600-650, 750-800, 800-850 and 900-950 ms, and strong (10<*K*<100) at 500-550 ms. These windows fall around the P100, on the descending limb of the P200, and throughout the late negativity; the P200 peak itself (200-300 ms) showed no evidence for an interaction (Figure 3**A**, shading). A list of all statistical contrasts is available in the Supplementary Table 5.

**Figure 3.**
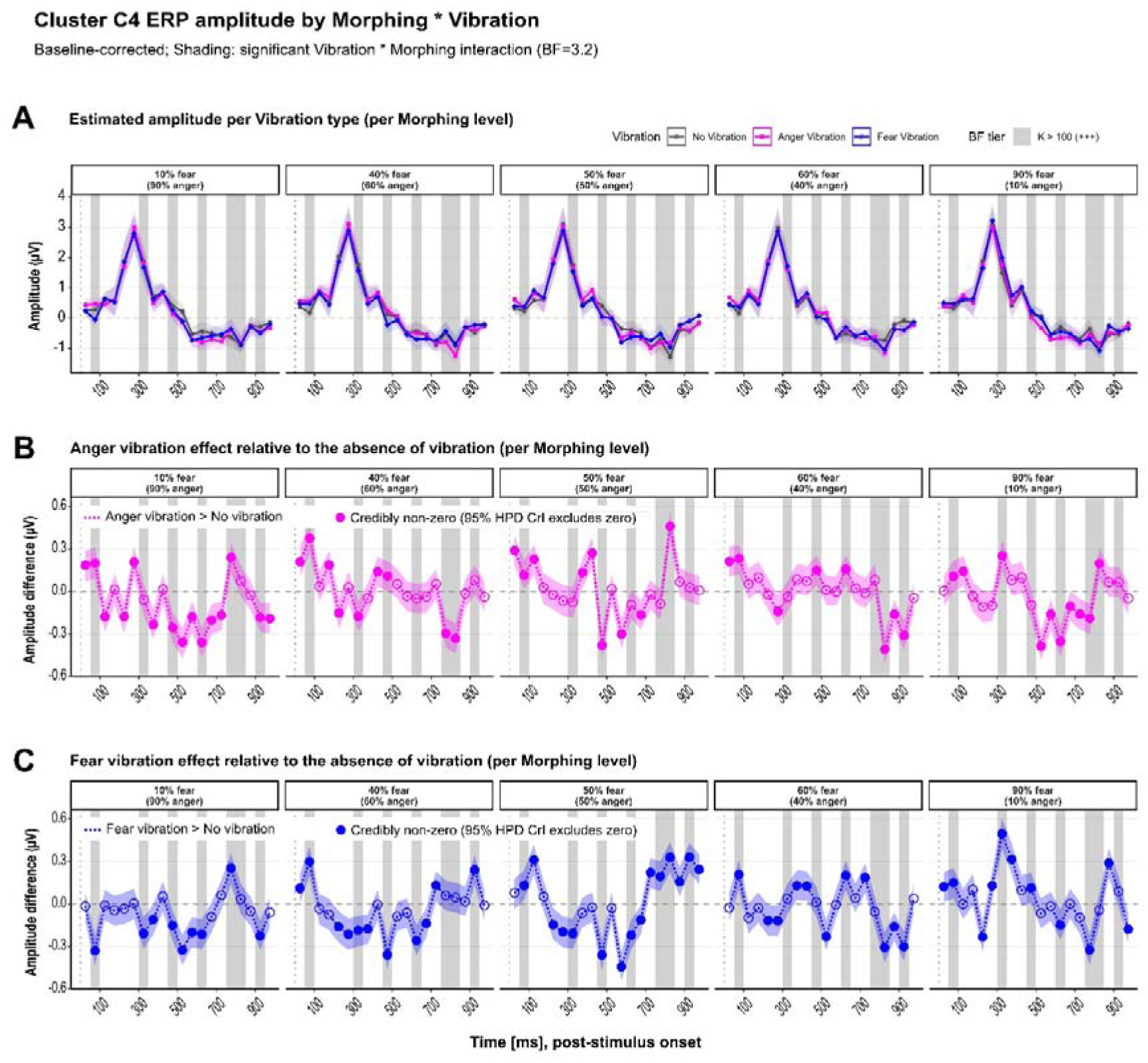
Cluster C4 frontocentral ERP amplitude as a function of Vibration type and Morphing level. (**A**) Estimated marginal mean amplitude (µV) per vibration condition across the five morphing levels, derived from Bayesian linear mixed models (brms; one model per 50ms timebin). Ribbons represent 95% highest posterior density credible intervals. Background shading indicates timebins where the Vibration * Morphing interaction reached decisive evidence (Bayes Factor *K*>100, +++; Jeffreys, 1998; Kass & Raftery, 1995). The dotted vertical line marks stimulus onset. (**B**) Amplitude difference (µV) between Anger Vibration and No Vibration conditions across morphing levels. Filled circles indicate timebins where the 95% credible interval of the pairwise contrast excludes zero (credibly non-zero); open circles indicate timebins where the credible interval includes zero. Ribbons represent 95% credible intervals of the contrast estimate. (**C**) Same as (**B**) for the Fear Vibration versus No Vibration contrast. Note that background shading in panels **B** and **C** reflects the significance of the overall Vibration * Morphing interaction (as in **A**), not of the individual pairwise contrasts shown. All models included participant and electrode as crossed random intercepts, and participant gender and voice gender as fixed covariates. Results are averaged over participant gender and voice actor gender.

Within these eight windows, we estimated the effect of each vibration relative to no vibration, at each morphing level (Figure 3**B**, **C**). Credible effects (95% credible interval, CrI, excluding zero) were present in 54 of 80 vibration * morphing level * window combinations. They were small (absolute difference=0.11–0.50 µV, against a P200 peak of about 3 µV) and their sign depended on window and morphing level. Negative difference indicates lower amplitude with vibration, i.e., a more negative-going signal.

At 50-100 ms, anger vibration increased amplitude at all five morphing levels (difference=0.11-0.37 µV; e.g., 40% fear: difference=0.37, 95% CrI [0.27, 0.48]), and fear vibration did so at 40-90% fear (difference=0.13-0.30 µV). For the least ambiguous, anger-dominant voice (10% fear), fear vibration instead decreased amplitude (difference=-0.33, 95% CrI [-0.44, -0.22]). On the descending limb of the P200 (300-350 ms), the effect of fear vibration changed sign along the continuum: it decreased amplitude at 10% (difference=-0.21, [-0.32, -0.10]), 40% (difference=-0.19, [-0.30, -0.09]) and 50% fear (difference=-0.21, [-0.32, -0.11]), had no credible effect at 60% fear, and increased amplitude at 90% fear (difference=0.50, [0.39, 0.60]). Anger vibration decreased amplitude at 40% fear (difference=-0.18, [-0.29, -0.07]) and increased it at 90% fear (difference=0.25, [0.14, 0.36]).

During the late negativity (450-950 ms), both vibrations decreased amplitude for the least ambiguous anger-dominant voice (10% fear) at 450-550, 600-650 and 900-950 ms (anger: difference=-0.18 to -0.36 µV; fear: difference=-0.15 to -0.33 µV) and increased it at 750-800 ms (anger: difference=0.24, [0.13, 0.34]; fear: difference=0.25, [0.14, 0.36]). For the most fearful voice (90% fear), anger vibration decreased amplitude at 500-550 and 600-650 ms (difference=-0.39, [-0.49, -0.28]; difference=-0.36, [-0.46, -0.25]). For ambiguous voices, effects reversed between windows. At 50% fear, anger and fear vibration both decreased amplitude at 450-500 ms (anger: difference=-0.38, [-0.48, -0.27]; fear: difference=-0.36, [-0.47, -0.25]) and increased it at 800-850 ms (anger: difference=0.46, [0.35, 0.56]; fear: difference=0.33, [0.22, 0.44]). At 60% fear, both vibrations decreased amplitude at 800-850 ms (anger: difference=-0.41, [-0.52, -0.30]; fear: difference=-0.31, [-0.41, -0.20]) and at 900-950 ms (anger: difference=-0.31, [-0.42, -0.21]; fear: difference=-0.30, [-0.40, -0.20]).

Credible effects occurred for the combinations that were behaviorally effective, namely fear vibration at 10-40% fear (5-7 of the 8 windows) and anger vibration at 50-60% fear (3-5 windows), but also for the remaining combinations, and anger and fear vibration often shifted amplitude in the same direction (Figure 3**B**, **C**).

#### 3.2.2 Source reconstruction

To localize these modulations, we reconstructed cortical sources with an LCMV beamformer and, for each of the 360 HCP-MMP1 ROIs and each of the eight 50-ms windows selected from the ERP results (75, 325, 475, 525, 625, 775, 825 and 925 ms), tested the effect of each vibration relative to no vibration within each morphing level (Figure 4; Supplementary Table 9). In total, 55 ROI-level effects reached significance after sequential GLMM analysis and survived correction (*p*FDR<.05): 20 for anger vibration (Figure 4**A**) and 35 for fear vibration (Figure 4**B**), 53 of them in the right hemisphere. *Z*-ratios refer to the contrast vibration minus no vibration and are positive when source amplitude was higher with vibration.

**Figure 4.**
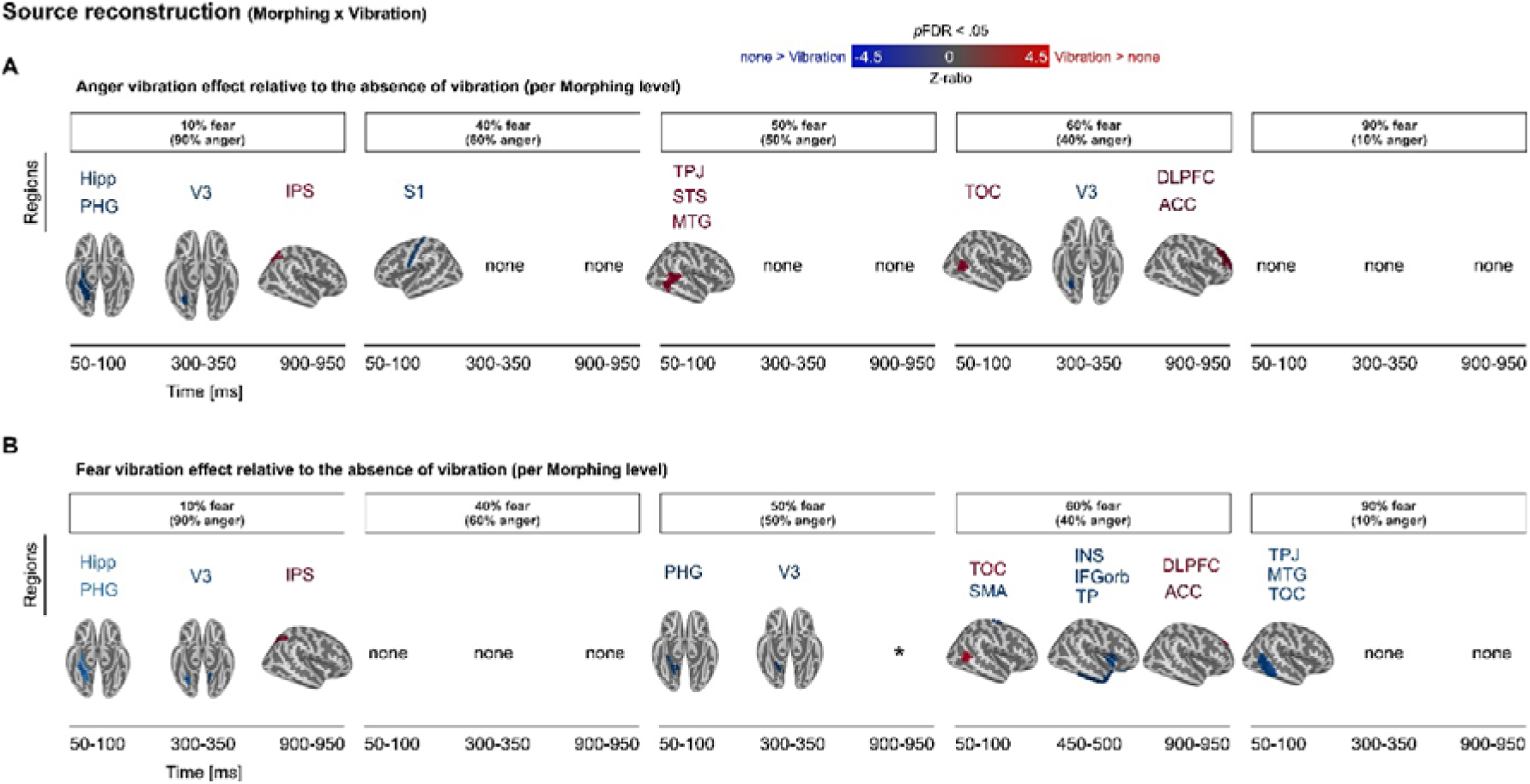
Source-level effects of vibration relative to no vibration, per morphing level. Sources were reconstructed from the 64-channel EEG with an LCMV beamformer (fsaverage template, 3-layer BEM; signal window 0-1 s, noise window -1 to 0 s) and averaged within the 360 parcels of the HCP-MMP1 atlas. For each ROI and 50-ms window, trial-level source amplitude was modelled with a Gaussian mixed model (Morphing × Vibration, random intercept for participant). The effect of Vibration was tested within each morphing level, using contrasts against the No-Vibration condition (emmeans). Outliers beyond 3 × IQR were excluded. Windows were chosen on the basis of the sensor-level ERP effects. Only ROIs surviving correction for multiple comparisons are shown (*p*FDR < .05). (**A**) Anger Vibration versus No Vibration. (**B**) Fear Vibration versus No Vibration. Colour indicates the z-ratio of the contrast: red, higher source amplitude with vibration than without (vibration > none); blue, higher without vibration (none > vibration); scale range ±4.5. Statistics were computed at ROI level and are projected onto the vertices of each parcel for display, so the extent of a coloured patch reflects parcel boundaries rather than sub-parcel localization. ACC, anterior cingulate cortex; Hipp, hippocampus; PHG, parahippocampal gyrus; V3, visual area 3; IPS, intraparietal sulcus; S1, primary somatosensory cortex; TPJ, temporo-parietal junction; STS, superior temporal sulcus; MTG, middle temporal gyrus; TOC, temporo-occipital cortex; DLPFC, dorsolateral prefrontal cortex; INS, insula; IFGorb, orbital inferior frontal gyrus; TP, temporal pole; SMA, supplementary motor area. ‘none’: no ROI survived correction. * medial ROI, not visible in these rendered views, see Supplementary Table 9.

At 50-100 ms, effects were found at every morphing level. For the least ambiguous anger-dominant voice (10% fear), both anger and fear vibration lowered source amplitude in right hippocampal, parahippocampal and ventromedial visual (V3) ROIs (anger: *z*=-3.30 to -3.68; fear: *z*=-2.18 to -2.76). At 40% fear, anger vibration lowered amplitude in left primary somatosensory area (*z*=-3.71). At 50% fear, anger vibration raised amplitude in right temporo-parietal junction (TPJ), and posterior temporal ROIs (*z*=3.44 and 3.35), whereas fear vibration lowered it in parahippocampal, ventromedial visual ROIs (*z*=-3.22 to -3.49). At 60% fear, both vibrations raised amplitude in right temporo-occipital cortex (anger: *z*=3.61; fear: *z*=2.75), and fear vibration lowered it in supplementary motor area (*z*=-3.47). At 90% fear, fear vibration lowered amplitude in right TPJ, posterior temporal cortex (*z*=-2.82 to -3.04).

At 300-350 ms, effects were confined to ventromedial visual and prostriate ROIs. Anger vibration lowered amplitude in right V3 at 10% and 60% fear (*z*=-3.47 for both). Fear vibration lowered it in right V3 (*z*=-2.73) and left prostriate cortex (*z*=-2.89) at 10% fear, and in right ventromedial visual area 1 (*z*=-3.43) at 50% fear. No ROI survived at 40% or 90% fear.

At 450-500 ms, the only surviving effect was for fear vibration at 60% fear, which lowered amplitude in right insular, orbitofrontal and anterior and medial temporal ROIs (*z*=-2.69 to -4.39). One further effect at 750-800 ms (fear vibration, 40% fear, right cingulate area p24, *z*=2.62) is listed in the Supplementary Table 9.

At 900-950 ms, effects were found in parietal, cingulate and prefrontal ROIs. At 10% fear, both vibrations raised amplitude in right intraparietal sulcus (IPS; anger: *z*=2.71 to 3.52; fear: *z*=2.28 to 3.23). At 50% fear, fear vibration raised amplitude in right medial parietal cortex (*z*=3.64). At 60% fear, anger vibration raised amplitude in dorsolateral prefrontal cortex (DLPFC; *z*=3.54 to 4.32) and anterior cingulate cortex (ACC; *z*=4.04 and 3.68), and fear vibration did so in the DLPFC (*z*=2.56) and ACC (*z*=2.23). No ROI survived at 40% or 90% fear.

Source-level effects were therefore right-lateralized and window-specific: ventral visual and medial temporal cortex early (50-350 ms), insular, orbitofrontal and anterior temporal cortex for fear vibration at 450-500 ms, and parietal, prefrontal and cingulate cortex late (900-950 ms). They included the insular, somatosensory, prefrontal, cingulate and supplementary motor regions anticipated in our source-level hypothesis, each at only a few combinations of morphing level and vibration. Within the windows shown in Figure 4, anger vibration had no surviving effect at 90% fear and fear vibration none at 40% fear. See Supplementary Table 9 for details.

## 4 Discussion

Throat vibrations changed how the listeners categorized emotional voices, in a way that depended on the morphing level. They also modulated fronto-central (C4) ERP amplitude in specific early and late windows and were associated with a distributed set of source effects, partly in the expected regions.

As mentioned, our study highlighted an impact of induced vibrations on the behavioral responses to morphed emotional voices, which is in line with our main hypothesis. Relative to no vibration, anger vibrations lowered the proportion of fear responses at 50% and 60% fear (OR 0.76 and 0.77), and fear vibrations raised it at 10% and 40% fear (OR 1.45 and 1.34). No effect was found for the clearest fear voices (90%). Each effect shifted categorization toward the emotion carried by the vibration, as expected if vibrotactile feedback is weighed during perception, and as motor stimulation biases speech perception (D’Ausilio et al., 2009). This is also coherent with studies on emotion-specific cross-modal bias (Lin & Ding, 2019; Niedenthal et al., 2001). The effect for the most ambiguous voices (50 and 60% fear) is consistent with the SIMS model (Niedenthal et al., 2010), in which bodily feedback weighs more for non-prototypical stimuli; the fear-vibration effect for anger-leaning voices is not predicted by it.

These results stand in apparent contrast to an earlier study from our group using a similar vibrotactile manipulation (Selosse et al., 2022), which found that congruent vibrations increased confusion rather than facilitating recognition, with no significant Vibration * Ambiguity interaction. We interpret this discrepancy as reflecting differences in design rather than in the underlying mechanism. That study manipulated ambiguity through acoustic intensity and used independent rating scales, which can register general disruption but not a directional bias. Morphing two emotions and forcing a binary choice, as that paper proposed, can reveal such a directional shift.

At the neural level, the frontocentral cluster C4 was the locus of an interaction between voice ambiguity and induced throat vibrations—both for early and late time-windows. At C4, the earliest effect (50 to 100 ms) was an amplitude increase with anger vibrations at all morphing levels and with fear vibrations at 40 to 90% fear, whereas fear vibrations reduced amplitude for the most anger-like voices (10% fear). In the literature, the P100 was linked to attention-driven interoceptive accuracy effects—auditory tone appearing in synchrony with heartbeat or with a delay (Mendoza-Medialdea & Ruiz-Padial, 2021). In cross-modal studies, multimodal integration of emotion recognition occurred very early, around 110ms post-stimulus onset (Pourtois et al., 2000). Furthermore, studies investigating interoception and somatosensory stimulation observed P300 modulations, particularly in the parietal area and such later effects therefore could be expected (Al et al., 2020; Auksztulewicz & Blankenburg, 2013; Has et al., 2021; Herbert et al., 2007; Pollatos et al., 2005).

We also observed the presence of a P200 component in this frontocentral area, consistent with emotional voice processing (Duville et al., 2022; Gädeke et al., 2013; Maltezou-Papastylianou et al., 2022; Paulmann et al., 2013; Paulmann & Uskul, 2017). After the P200 peak, around 300 to 350 ms, fear vibrations lowered amplitude for anger-leaning and ambiguous voices (10 to 50% fear) and raised it for the clearest fear voices (90%); anger vibrations showed a similar pattern at 40 and 90% fear. These results are in line with above-mentioned findings on multimodal integration and interoception, but in our results they occurred after, not at, the peak. These effects show that vibrations are registered early and again around 300 ms; they do not by themselves show that these modulations underlie the behavioral shift, which we did not test. Moreover, the effect of fear vibrations for ambiguous voice categorization only is also supported by the SIMS model (Niedenthal, 2007; Niedenthal et al., 2010).

Later, from 450 to 950 ms, anger and fear vibrations often shifted amplitude in the same direction (mostly a decrease early in this range, an increase around 750 to 850 ms) at several morphing levels, including the most ambiguous (50 and 60% fear). These results recall the N400 component observed in frontal, parietal and frontocentral areas during emotional voice processing (Föcker & Röder, 2019; Maltezou-Papastylianou et al., 2022; Tarai & Srinivasan, 2019). They are also reminiscent of late positive modulations, linked to emotional ‘meaning’ (Wei et al., 2022; Zora et al., 2020), prosody expectancy violation (Astésano et al., 2004; Brattico et al., 2006; Paulmann et al., 2012) and arousal (Paulmann et al., 2013), but our design cannot assign them to a specific process. Nevertheless, this echoes once again with the SIMS model—embodied simulation is engaged when the emotional stimulus is non-prototypical (Niedenthal et al., 2010). Here, the vibrations were integrated for ambiguous voices (voices with 50% and 60% fear), as prototypical emotional voice processing would not be enough to identify the emotion. At the interoception and proprioception levels, these results globally show that vibrations are integrated during emotional processing at multiple stages, dynamically and throughout the time course of an event, similarly to the multimodal integration shown in audiovisual studies (Föcker & Röder, 2019; Klasen et al., 2012).

ERP source reconstruction revealed source modulations in several key regions. Some of these regions are involved in vibrotactile processing: the DLPFC in working memory (Burton et al., 2010), and also others in vibratory processing, crucially in vibration rate or pattern discrimination (Christensen et al., 2007; Coghill et al., 1994; Golaszewski et al., 2006; Hernández et al., 2002; Lenoir et al., 2017; Li Hegner et al., 2010; Macar et al., 2002; Romo et al., 2004; Woolgar & Zopf, 2017; Wu et al., 2018). Concerning more internal feedback or interoception, we observed source modulations in the insula, known to be crucially involved in interoceptive processing (Adolfi et al., 2017; Casals-Gutierrez & Abbey, 2020) and interoceptive awareness (Herbert et al., 2007; Longarzo et al., 2021; Wilson-Mendenhall et al., 2019). The modulation of the primary somatosensory cortex might be related to proprioceptive information processing while modulations in the OFC were also reported in interoception studies (Singer & Klimecki, 2014; Stern et al., 2017).

Regarding emotional prosody perception, and among other regions, we observed source modulations in the DLPFC (prosodic working memory; Mitchell, 2007), superior parietal cortex (emotional prosody judgement; Bach et al., 2008), as well as in other regions also involved in emotional prosody processing (IPS, SMA; Alba-Ferrara et al., 2011; Belyk & Brown, 2014; Kotz et al., 2003). OFC modulations fit well with higher-level processing of emotional prosody (Wildgruber et al., 2006)—especially when ambiguous (Grandjean, 2021), or for accurately categorizing human voice (Ceravolo et al., 2023). Source modulations in the insula may contribute to external emotional stimuli integration (vibrotactile feedback) and emotional experiences with interoceptive signals (Adolfi et al., 2017; Nguyen et al., 2016; Zaki et al., 2012). Finally, above-mentioned modulations—superior parietal cortex, IPS, supplementary motor area, primary somatosensory cortex—are in line with multimodal integration for auditory, visual, sensorimotor and tactile stimuli (Baker et al., 2018; Culham & Valyear, 2006; de Borst & de Gelder, 2017; Gentile et al., 2011; Kreifelts et al., 2007; Parker et al., 2020). Source modulation locations for the Vibration and Morphing factors are therefore coherent with our previous functional MRI results on neural correlates of emitted, measured vocal tract vibrations during emotional expression (Selosse et al., 2025). Combined with the present data, they help draw a clearer picture of embodied accounts in emotional voice signals. Source tests were however exploratory and based on C4 cluster ERP time windows.

More generally, the voice is a bodily signal before it is an acoustic one: producing a vocal emotion changes laryngeal and vocal tract vibrations that the speaker feels, and these self-generated vibrations are available as proprioceptive and interoceptive information during production (Selosse et al., 2025). Our results suggest that the same channel can be re-used during perception. This is in line with work showing that listening to speech engages motor and premotor regions (Fadiga et al., 2002; Watkins et al., 2003), and with the single SMA effect we found, which would fit a link between hearing and producing a vocal emotion. Vibrations applied to the throat of a listener shifted how morphed voices were categorized, each vibration pulling the response toward its own emotion, and this occurred at a range of ambiguity levels except the clearest fear voices. This fits accounts in which perceiving an emotion partly re-enacts it in the perceiver’s body (Niedenthal, 2007; Niedenthal et al., 2010), and extends them from facial mimicry and posture to a vibrotactile component of the voice itself. The ambiguity gradient is a useful way to see this: bodily feedback should weigh most when the acoustic evidence alone is insufficient, and in our data it did bias the responses where voices were mixed (50 and 60% fear), but also for anger-leaning voices (10 and 40% fear), suggesting that the relevant limit is not only the absence of evidence but also its relative weight against the vibration. We cannot say whether the vibrations were integrated with the acoustic signal or acted as a response bias; the ERP and source results, being small and window-specific, do not settle this.

Together, these results indicate that throat vibrations bias vocal emotion categorization, each vibration pulling responses toward its own emotion, and that this is accompanied by small, time-specific ERP changes and a distributed set of source effects. Whether vibrotactile feedback is necessary for vocal emotion perception remains an open question.

### 4.1 Limitations and perspectives

Firstly, our sample consists mainly of female participants, despite known gender differences in emotional prosody recognition (Imaizumi et al., 2004; Lambrecht et al., 2014) and facial mimicry (Korb et al., 2015; Niedenthal et al., 2012). Second, it would be necessary to replicate the observed effects with different emotions to generalize our results, although morphing positive and negative emotions can be problematic. Third, induced vibrations should be those of the participants’ own expressions for improved ecological validity—and better consideration of interindividual differences. Fourth, vibrations were induced on the skin, therefore more subtle internal tissue vibrations encoded by different mechano-receptors may not be triggered. Further research is needed to clarify the relationships between actual vibrations created by the vocal cords and those induced on the skin level. Also, we tested perception only, so whether the same vibrations also support vocal emotion production remains open. Finally, future studies would benefit from the inclusion of measures of bodily self-consciousness or interoceptive awareness. In fact, interindividual variability in this dimension probably impacts subsequent data as the feedback mechanism highlighted in the present work might highly depend on such sensitivity. The effects could therefore scale with interoceptive abilities.

## Data availability statement

Data and codes can be downloaded from the open, FAIR-compliant Yareta repository under the following permanent link: https://doi.org/10.26037/yareta:eu3ihgdnqzdj3g4ayn2xzti6wq.

## References

Adolfi, F., Couto, B., Richter, F., Decety, J., Lopez, J., Sigman, M., & Ibáñez, A. (2017). Convergence of interoception, emotion, and social cognition: a twofold fMRI meta-analysis and lesion approach. Cortex, 88, 124--142.

Al, E., Iliopoulos, F., Forschack, N., Nierhaus, T., Grund, M., Motyka, P., & Villringer, A. (2020). Heart–brain interactions shape somatosensory perception and evoked potentials. Proceedings of the National Academy of Sciences, 117(19), 10575--10584.

Alba-Ferrara, L., Hausmann, M., Mitchell, R. L., & Weis, S. (2011). The Neural Correlates of Emotional Prosody Comprehension: Disentangling Simple from Complex Emotion. PLoS ONE, 6(12), 28701. 10.1371/journal.pone.0028701

Astésano, C., Besson, M., & Alter, K. (2004). Brain potentials during semantic and prosodic processing in French. Cognitive Brain Research, 18(2), 172--184.

Auksztulewicz, R., & Blankenburg, F. (2013). Subjective rating of weak tactile stimuli is parametrically encoded in event-related potentials. Journal of Neuroscience, 33(29), 11878--11887.

Bach, D. R., Grandjean, D., Sander, D., Herdener, M., Strik, W. K., & Seifritz, E. (2008). The effect of appraisal level on processing of emotional prosody in meaningless speech. NeuroImage, 42(2), 919--927. 10.1016/j.neuroimage.2008.05.034

Baker, C. M., Burks, J. D., Briggs, R. G., Conner, A. K., Glenn, C. A., Taylor, K. N., Sali, G., McCoy, T. M., Battiste, J. D., O’Donoghue, D. L., & Sughrue, M. E. (2018). A Connectomic Atlas of the Human Cerebrum—Chapter 7: The Lateral Parietal Lobe. Operative Neurosurgery, 15(suppl_1), 295--349. 10.1093/ons/opy261

Bates, D., Maechler, M., Bolker, B., Walker, S., Christensen, R. H. B., Singmann, H., Dai, B., Grothendieck, G., Green, P., & Bolker, M. B. (2015). Package ‘lme4’. convergence, 12(1), 2.

Belin, P., Fillion-Bilodeau, S., & Gosselin, F. (2008). The Montreal Affective Voices: A validated set of nonverbal affect bursts for research on auditory affective processing. Behavior research methods, 40(2), 531--539.

Belyk, M., & Brown, S. (2014). Perception of affective and linguistic prosody: An ALE meta-analysis of neuroimaging studies. Social Cognitive and Affective Neuroscience, 9(9), 1395--1403. 10.1093/scan/nst124

Blairy, S., Herrera, P., & Hess, U. (1999). Mimicry and the judgment of emotional facial expressions. Journal of Nonverbal Behavior, 23, 5--41.

Borgomaneri, S., Bolloni, C., Sessa, P., & Avenanti, A. (2020). Blocking facial mimicry affects recognition of facial and body expressions. PLoS ONE, 15(2), e0229364.

Brainard, D. H. (1997). The psychophysics toolbox. Spatial Vision, 10(4), 433--436. 10.1163/156856897x00357

Brattico, E., Tervaniemi, M., Näätänen, R., & Peretz, I. (2006). Musical scale properties are automatically processed in the human auditory cortex. Brain Research, 1117(1), 162--174.

Brosch, T., Grandjean, D., Sander, D., & Scherer, K. R. (2008). Behold the voice of wrath: Cross-modal modulation of visual attention by anger prosody. Cognition, 106(3), 1497--1503.

Brosch, T., Sander, D., Pourtois, G., & Scherer, K. R. (2008). Beyond fear: Rapid spatial orienting toward positive emotional stimuli. Psychological science, 19(4), 362--370.

Brunswik, E. (1956). *Perception and the representative design of psychological experiments*. University of California Press.

Bürkner, P.-C. (2017). brms: An R package for Bayesian multilevel models using Stan. Journal of statistical software, 80, 1–28.

Burleson, M. H., & Quigley, K. S. (2021). Social interoception and social allostasis through touch: legacy of the somatovisceral afference model of emotion. Social neuroscience, 16(1), 92--102.

Burton, H., Sinclair, R. J., & Dixit, S. (2010). Working memory for vibrotactile frequencies: Comparison of cortical activity in blind and sighted individuals. Human Brain Mapping, 31(11), 1686--1701. 10.1002/hbm.20966

Carminati, M., Fiori-Duharcourt, N., & Isel, F. (2018). Neurophysiological differentiation between preattentive and attentive processing of emotional expressions on French vowels. Biological psychology, 132, 55--63.

Casals-Gutierrez, S., & Abbey, H. (2020). Interoception, mindfulness and touch: A meta-review of functional MRI studies. International Journal of Osteopathic Medicine, 35, 22--33.

Ceravolo, L., Debracque, C., Pool, E., Gruber, T., & Grandjean, D. (2023). Frontal mechanisms underlying primate calls recognition by humans. Cerebral Cortex Communications, 4(4), tgad019.

Ceravolo, L., Frühholz, S., Pierce, J., Grandjean, D., & Péron, J. (2021). Basal ganglia and cerebellum contributions to vocal emotion processing as revealed by high-resolution fMRI. Scientific reports, 11(1), 10645.

Ceravolo, L., Moisa, M., Grandjean, D., Ruff, C., & Frühholz, S. (2026). Behavioral and neural aftermath of right inferior frontal cortex disruption on ambiguous vocal emotion decisional processes. Scientific reports. 10.1038/s41598-026-39668-0

Chartrand, T. L., & Bargh, J. A. (1999). The chameleon effect: The perception–behavior link and social interaction. Journal of personality and social psychology, 76(6), 893.

Christensen, M. S., Lundbye-Jensen, J., Geertsen, S. S., Petersen, T. H., Paulson, O. B., & Nielsen, J. B. (2007). Premotor cortex modulates somatosensory cortex during voluntary movements without proprioceptive feedback. Nature Neuroscience, 10(4), 417--419. 10.1038/nn1873

Coghill, R., Talbot, J., Evans, A., Meyer, E., Gjedde, A., Bushnell, M., & Duncan, G. (1994). Distributed processing of pain and vibration by the human brain. The Journal of Neuroscience, 14(7), 4095--4108. 10.1523/jneurosci.14-07-04095.1994

Couto, B., Adolfi, F., Velasquez, M., Mesow, M., Feinstein, J., Canales-Johnson, A., & Ibanez, A. (2015). Heart evoked potential triggers brain responses to natural affective scenes: a preliminary study. Autonomic Neuroscience, 193, 132--137.

Critchley, H. D., & Garfinkel, S. N. (2017). Interoception and emotion. Current opinion in psychology, 17, 7--14.

Culham, J. C., & Valyear, K. F. (2006). Human parietal cortex in action. Current Opinion in Neurobiology, 16(2), 205--212. 10.1016/j.conb.2006.03.005

D’Ausilio, A., Pulvermüller, F., Salmas, P., Bufalari, I., Begliomini, C., & Fadiga, L. (2009). The motor somatotopy of speech perception. Current Biology, 19(5), 381--385.

de Borst, A. W., & de Gelder, B. (2017). fMRI-based multivariate pattern analyses reveal imagery modality and imagery content specific representations in primary somatosensory, motor and auditory cortices. Cerebral Cortex, 27(8), 3994--4009. 10.1093/cercor/bhw211

Dimberg, U. (1982). Facial reactions to facial expressions. Psychophysiology, 19(6), 643--647.

Drimalla, H., Landwehr, N., Hess, U., & Dziobek, I. (2019). From face to face: the contribution of facial mimicry to cognitive and emotional empathy. Cognition and Emotion.

Duville, M. M., Alonso-Valerdi, L. M., & Ibarra-Zarate, D. I. (2022). Neuronal and behavioral affective perceptions of human and naturalness-reduced emotional prosodies. Frontiers in Computational Neuroscience, 16, 1022787.

Engelen, T., Solcà, M., & Tallon-Baudry, C. (2023). Interoceptive rhythms in the brain. Nature Neuroscience, 26(10), 1670--1684.

Fadiga, L., Craighero, L., Buccino, G., & Rizzolatti, G. (2002). Speech listening specifically modulates the excitability of tongue muscles: a TMS study. European Journal of Neuroscience, 15(2), 399--402.

Faul, F., Erdfelder, E., Lang, A. G., & Buchner, A. (2007). G* Power 3: A flexible statistical power analysis program for the social, behavioral, and biomedical sciences. Behavior research methods, 39(2), 175--191.

Filippa, M., Grandjean, D., & Ceravolo, L. (2026). Toward An Embodied Feedback Model of Vocal Emotion Production: Linking Subjective Perception, Objective Vibratory Dynamics, And Interoception Sensitivity [Preprint]. SSRN. 10.2139/ssrn.7198221

Föcker, J., & Röder, B. (2019). Event-related potentials reveal evidence for late integration of emotional prosody and facial expression in dynamic stimuli: An ERP study. Multisensory Research, 32(6), 473--497.

Gädeke, J. C., Föcker, J., & Röder, B. (2013). Is the processing of affective prosody influenced by spatial attention? An ERP study. BMC neuroscience, 14(1), 1--15.

Gentile, G., Petkova, V. I., & Ehrsson, H. H. (2011). Integration of Visual and Tactile Signals From the Hand in the Human Brain: An fMRI Study. Journal of Neurophysiology, 105(2), 910--922. 10.1152/jn.00840.2010

Gibson, J. (2019). Mindfulness, interoception, and the body: A contemporary perspective. Frontiers in Psychology, 10.

Glasser, M. F., Coalson, T. S., Robinson, E. C., Hacker, C. D., Harwell, J., & Yacoub, E. (2016). A multi-modal parcellation of human cerebral cortex. Nature, 1--11. 10.1038/nature18933

Golaszewski, S. M., Siedentopf, C. M., Koppelstaetter, F., Fend, M., Ischebeck, A., Gonzalez-Felipe, V., Haala, I., Struhal, W., Mottaghy, F. M., Gallasch, E., Felber, S. R., & Gerstenbrand, F. (2006). Human brain structures related to plantar vibrotactile stimulation: A functional magnetic resonance imaging study. NeuroImage, 29(3), 923--929. 10.1016/j.neuroimage.2005.08.052

Grandjean, D. (2021). Brain networks of emotional prosody processing. Emotion Review, 13(1), 34--43.

Grandjean, D., Bänziger, T., & Scherer, K. R. (2006). Intonation as an interface between language and affect. Progress in brain research, 156, 235--247.

Has, N. A., Jamaludin, M. N., Rejab, S. M., Ahmad, Z., Zainuddin, N. F., Salam, A. H. A., & As’ari, M. A. (2021). P300 Somatosensory Validation of Vibrotactile Haptic Feedback for Upper Limb Prosthesis. 2021 IEEE National Biomedical Engineering Conference (NBEC),

Hawk, S. T., Fischer, A. H., & van Kleef, G. A. (2012). Face the noise: embodied responses to nonverbal vocalizations of discrete emotions. Journal of personality and social psychology, 102(4), 796.

Herbert, B. M., Pollatos, O., & Schandry, R. (2007). Interoceptive sensitivity and emotion processing: an EEG study. International Journal of Psychophysiology, 65(3), 214--227.

Hernández, A., Zainos, A., & Romo, R. (2002). Temporal Evolution of a Decision-Making Process in Medial Premotor Cortex. Neuron, 33(6), 959--972. 10.1016/s0896-6273(02)00613-x

Hess, U., & Blairy, S. (2001). Facial mimicry and emotional contagion to dynamic emotional facial expressions and their influence on decoding accuracy. International Journal of Psychophysiology, 40(2), 129--141.

Hess, U., & Fischer, A. (2014). Emotional mimicry: Why and when we mimic emotions. Social and personality psychology compass, 8(2), 45--57.

Holland, A. C., O’Connell, G., & Dziobek, I. (2021). Facial mimicry, empathy, and emotion recognition: a meta-analysis of correlations. Cognition and Emotion, 35(1), 150--168.

Hyniewska, S., & Sato, W. (2015). Facial feedback affects valence judgments of dynamic and static emotional expressions. Frontiers in Psychology, 6, 291.

Imaizumi, S., Homma, M., Ozawa, Y., Maruishi, M., & Muranaka, H. (2004). Gender differences in emotional prosody processing—An fMRI study. Psychologia, 47(2), 113--124.

Jeffreys, H. (1998). The theory of probability. OuP Oxford.

Kass, R. E., & Raftery, A. E. (1995). Bayes factors. Journal of the american statistical association, 90(430), 773–795.

Kawahara, H., & Matsui, H. (2003). Auditory morphing based on an elastic perceptual distance metric in an interference-free time-frequency representation. 2003 IEEE International Conference on Acoustics, Speech, and Signal Processing,

Khalsa, S. S., Adolphs, R., Cameron, O. G., Critchley, H. D., Davenport, P. W., Feinstein, J. S., & Zucker, N. (2018). Interoception and mental health: a roadmap. Biological psychiatry: cognitive neuroscience and neuroimaging, 3(6), 501--513.

Klasen, M., Chen, Y. H., & Mathiak, K. (2012). Multisensory emotions: perception, combination and underlying neural processes. Reviews in the Neurosciences, 23(4), 381--392.

Kleiner, M., Brainard, D., & Pelli, D. (2007). What’s new in Psychtoolbox-3. In.

Korb, S., Grandjean, D., & Scherer, K. R. (2010). Timing and voluntary suppression of facial mimicry to smiling faces in a Go/NoGo task—An EMG study. Biological psychology, 85(2), 347--349.

Korb, S., Malsert, J., Rochas, V., Rihs, T. A., Rieger, S. W., Schwab, S., & Grandjean, D. (2015). Gender differences in the neural network of facial mimicry of smiles–An rTMS study. Cortex, 70, 101--114.

Kotz, S. A., Meyer, M., Alter, K., Besson, M., von Cramon, D. Y., & Friederici, A. D. (2003). On the lateralization of emotional prosody: An event-related functional MR investigation. Brain and Language, 86(3), 366--376. 10.1016/s0093-934x(02)00532-1

Kreifelts, B., Ethofer, T., Grodd, W., Erb, M., & Wildgruber, D. (2007). Audiovisual integration of emotional signals in voice and face: An event-related fMRI study. NeuroImage, 37(4), 1445--1456. 10.1016/j.neuroimage.2007.06.020

Künecke, J., Wilhelm, O., & Sommer, W. (2017). Emotion recognition in nonverbal face-to-face communication. Journal of Nonverbal Behavior, 41, 221--238.

Lambrecht, L., Kreifelts, B., & Wildgruber, D. (2014). Gender differences in emotion recognition: Impact of sensory modality and emotional category. Cognition & Emotion, 28(3), 452--469.

Laukka, P., Juslin, P., & Bresin, R. (2005). A dimensional approach to vocal expression of emotion. Cognition & Emotion, 19(5), 633--653.

Lenoir, C., Huang, G., Vandermeeren, Y., Hatem, S. M., & Mouraux, A. (2017). Human primary somatosensory cortex is differentially involved in vibrotaction and nociception. Journal of Neurophysiology, 118(1), 317--330. 10.1152/jn.00615.2016

Lenth, R. V. (2021). emmeans: Estimated Marginal Means, aka Least-Squares Means. In.

Li Hegner, Y., Lee, Y., Grodd, W., & Braun, C. (2010). Comparing Tactile Pattern and Vibrotactile Frequency Discrimination: A Human fMRI Study. Journal of Neurophysiology, 103(6), 3115--3122. 10.1152/jn.00940.2009

Lin, Y., & Ding, H. (2019). Multisensory integration of emotions in a face-prosody-semantics Stroop task. In NeuroManagement and Intelligent Computing Method on Multimodal Interaction (pp. 1--5).

Lindín, M., Zurrón, M., & Díaz, F. (2004). Changes in P300 amplitude during an active standard auditory oddball task. Biological psychology, 66(2), 153--167.

Longarzo, M., Mele, G., Alfano, V., Salvatore, M., & Cavaliere, C. (2021). Gender Brain Structural Differences and Interoception. Frontiers in Neuroscience, 14, 586860. 10.3389/fnins.2020.586860

Lylykangas, J., Surakka, V., Rantala, J., Raisamo, J., Raisamo, R., & Tuulari, E. (2009). Vibrotactile information for intuitive speed regulation. In People and Computers XXIII Celebrating People and Technology (pp. 112--119).

Macar, F., Lejeune, H., Bonnet, M., Ferrara, A., Pouthas, V., Vidal, F., & Maquet, P. (2002). Activation of the supplementary motor area and of attentional networks during temporal processing. Experimental Brain Research, 142(4), 475--485.

Maltezou-Papastylianou, C., Russo, R., Wallace, D., Harmsworth, C., & Paulmann, S. (2022). Different stages of emotional prosody processing in healthy ageing–evidence from behavioural responses, ERPs, tDCS, and tRNS. PLoS ONE, 17(7), e0270934.

Mendoza-Medialdea, M. T., & Ruiz-Padial, E. (2021). Understanding the capture of exogenous attention by disgusting and fearful stimuli: the role of interoceptive accuracy. International Journal of Psychophysiology, 161, 53--63.

Mitchell, R. L. C. (2007). fMRI delineation of working memory for emotional prosody in the brain: Commonalities with the lexico-semantic emotion network. NeuroImage, 36(3), 1015--1025. 10.1016/j.neuroimage.2007.03.016

Moody, E. J., & McIntosh, D. N. (2011). Mimicry of dynamic emotional and motor-only stimuli. Social Psychological and Personality Science, 2(6), 679--686.

Moody, E. J., McIntosh, D. N., Mann, L. J., & Weisser, K. R. (2007). More than mere mimicry? The influence of emotion on rapid facial reactions to faces. Emotion, 7(2), 447.

Munger, J. B., & Thomson, S. L. (2008). Frequency response of the skin on the head and neck during production of selected speech sounds. The Journal of the Acoustical Society of America, 124(6), 4001--4012.

Nguyen, V. T., Breakspear, M., Hu, X., & Guo, C. C. (2016). The integration of the internal and external milieu in the insula during dynamic emotional experiences. NeuroImage, 124, 455--463. 10.1016/j.neuroimage.2015.08.078

Niedenthal, P. M. (2007). Embodying Emotion. Science, 316(5827), 1002--1005.

Niedenthal, P. M., Augustinova, M., Rychlowska, M., Droit-Volet, S., Zinner, L., Knafo, A., & Brauer, M. (2012). Negative relations between pacifier use and emotional competence. Basic and Applied Social Psychology, 34(5), 387--394.

Niedenthal, P. M., Brauer, M., Halberstadt, J. B., & Innes-Ker, Å. H. (2001). When did her smile drop? Facial mimicry and the influences of emotional state on the detection of change in emotional expression. Cognition & Emotion, 15(6), 853--864.

Niedenthal, P. M., Mermillod, M., Maringer, M., & Hess, U. (2010). The Simulation of Smiles (SIMS) model: Embodied simulation and the meaning of facial expression. Behavioral and brain sciences, 33(6), 417.

Nolan, M., Madden, B., & Burke, E. (2009). Accelerometer based measurement for the mapping of neck surface vibrations during vocalized speech. Annual International Conference of the IEEE Engineering in Medicine and Biology Society,

Oostenveld, R., Fries, P., Maris, E., & Schoffelen, J.-M. (2011). FieldTrip: open source software for advanced analysis of MEG, EEG, and invasive electrophysiological data. Computational intelligence and neuroscience, 2011(1), 156869.

Paquette, S., Rigoulot, S., Grunewald, K., & Lehmann, A. (2020). Temporal decoding of vocal and musical emotions: Same code, different timecourse? Brain Research, 146887.

Parker, A. G., Briggs, R. G., Conner, A. K., O’Neal, C. M., Bonney, P. A., Maxwell, B. D., Baker, C. M., Burks, J. D., Sali, G., & Sughrue, M. E. (2020). Parcellation-based tractographic modeling of the ventral attention network. Journal of the Neurological Sciences, 408, 116548.

Paulmann, S., Bleichner, M., & Kotz, S. A. (2013). Valence, arousal, and task effects in emotional prosody processing. Frontiers in Psychology, 4, 345.

Paulmann, S., Jessen, S., & Kotz, S. A. (2012). It’s special the way you say it: An ERP investigation on the temporal dynamics of two types of prosody. Neuropsychologia, 50(7), 1609--1620.

Paulmann, S., & Kotz, S. A. (2008). An ERP investigation on the temporal dynamics of emotional prosody and emotional semantics in pseudo-and lexical-sentence context. Brain and Language, 105(1), 59--69.

Paulmann, S., & Uskul, A. K. (2017). Early and late brain signatures of emotional prosody among individuals with high versus low power. Psychophysiology, 54(4), 555--565.

Pell, M. D., & Kotz, S. A. (2021). Comment: The next frontier: Prosody research gets interpersonal. Emotion Review, 13(1), 51--56.

Pelli, D. G. (1997). The videotoolbox software for visual psychophysics: Transforming numbers into movies. Spatial Vision, 10(4), 437--442.

Peron, J., Frühholz, S., Ceravolo, L., & Grandjean, D. (2016). Structural and functional connectivity of the subthalamic nucleus during vocal emotion decoding. Social Cognitive and Affective Neuroscience, 11(2), 349–356.

Philip, L., Martin, J. C., & Clavel, C. (2018). Rapid facial reactions in response to facial expressions of emotion displayed by real versus virtual faces. i-Perception, 9(4), 2041669518786527.

Pollatos, O., Kirsch, W., & Schandry, R. (2005). On the relationship between interoceptive awareness, emotional experience, and brain processes. Cognitive Brain Research, 25(3), 948--962.

Pourtois, G., de Gelder, B., Vroomen, J., Rossion, B., & Crommelinck, M. (2000). The time-course of intermodal binding between seeing and hearing affective information. Neuroreport, 11(6), 1329--1333.

Proverbio, A. M., Benedetto, F., & Guazzone, M. (2020). Shared neural mechanisms for processing emotions in music and vocalizations. European Journal of Neuroscience, 51(9), 1987--2007.

Risberg, A., & Lubker, J. (1978). Prosody and speechreading. Speech Transmission Laboratory Quarterly Progress Report and Status Report, 4, 1--16.

Romo, R., Hernández, A., & Zainos, A. (2004). Neuronal Correlates of a Perceptual Decision in Ventral Premotor Cortex. Neuron, 41(1), 165--173. 10.1016/s0896-6273(03)00817-1

Salamone, P. C., Legaz, A., Sedeño, L., Moguilner, S., Fraile-Vazquez, M., Campo, C. G., & Ibañez, A. (2021). Interoception primes emotional processing: multimodal evidence from neurodegeneration. Journal of Neuroscience, 41(19), 4276--4292.

Scherer, K. R. (2003). Vocal communication of emotion: A review of research paradigms. Speech communication, 40(1-2), 227--256.

Scherer, K. R., & Oshinsky, J. S. (1977). Cue utilization in emotion attribution from auditory stimuli. Motivation and Emotion, 1(4), 331--346.

Schirmer, A., & Kotz, S. A. (2006). Beyond the right hemisphere: brain mechanisms mediating vocal emotional processing. Trends in cognitive sciences, 10(1), 24--30.

Schneider, K. G., Hempel, R. J., & Lynch, T. R. (2013). That "poker face" just might lose you the game! The impact of expressive suppression and mimicry on sensitivity to facial expressions of emotion. Emotion, 13(5), 852.

Selosse, G., Grandjean, D., & Ceravolo, L. (2022). Influence of bodily resonances on emotional prosody perception. Frontiers in Psychology, 13, 1061930.

Selosse, G., Grandjean, D., & Ceravolo, L. (2025). Neural correlates of embodied and vibratory mechanisms associated with emotional prosody production. Social Cognitive and Affective Neuroscience, 20(1), nsaf084.

Singer, T., & Klimecki, O. M. (2014). Empathy and compassion. Current Biology, 24(18), 875--878.

Stel, M., & van Knippenberg, A. (2008). The role of facial mimicry in the recognition of affect. Psychological science, 19(10), 984.

Stern, E. R., Grimaldi, S. J., Muratore, A., Murrough, J., Leibu, E., Fleysher, L., & Burdick, K. E. (2017). Neural correlates of interoception: Effects of interoceptive focus and relationship to dimensional measures of body awareness. Human Brain Mapping, 38(12), 6068--6082.

Sundberg, J. (1992). Phonatory vibrations in singers: A critical review. Music Perception: An Interdisciplinary Journal, 9(3), 361--381.

Švec, J. G., Titze, I. R., & Popolo, P. S. (2005). Estimation of sound pressure levels of voiced speech from skin vibration of the neck. The Journal of the Acoustical Society of America, 117(3), 1386--1394.

Tarai, S., & Srinivasan, N. (2019). Emotional prosody Stroop effect in Hindi: an event related potential study. In Progress in brain research (Vol. 247, pp. 193--217).

Thomas, S. J., Johnstone, S. J., & Gonsalvez, C. J. (2007). Event-related potentials during an emotional Stroop task. International Journal of Psychophysiology, 63(3), 221--231.

Tsolaki, A., Kosmidou, V., Hadjileontiadis, L., Kompatsiaris, I. Y., & Tsolaki, M. (2015). Brain source localization of MMN, P300 and N400: Aging and gender differences. Brain Research, 32--49.

Tuuri, K., Eerola, T., & Pirhonen, A. (2010). Leaping across modalities: Speed regulation messages in audio and tactile domains. International Workshop on Haptic and Audio Interaction Design,

Venables, W. N., & Smith, D. M. (2003). The R development core team. An Introduction to R, Version, 1(0).

Watkins, K. E., Strafella, A. P., & Paus, T. (2003). Seeing and hearing speech excites the motor system involved in speech production. Neuropsychologia, 41(8), 989--994.

Wei, H., He, Y., Kauschke, C., Scharinger, M., & Domahs, U. (2022). An EEG-study on L2 categorization of emotional prosody in German. Speech Prosody,

Wildgruber, D., Ackermann, H., Kreifelts, B., & Ethofer, T. (2006). Cerebral processing of linguistic and emotional prosody: fMRI studies. In Progress in brain research (Vol. 156, pp. 249--268). 10.1016/s0079-6123(06)56013-3

Wilson-Mendenhall, C. D., Henriques, A., Barsalou, L. W., & Barrett, L. F. (2019). Primary Interoceptive Cortex Activity during Simulated Experiences of the Body. Journal of Cognitive Neuroscience, 31(2), 221--235. 10.1162/jocn_a_01346

Wilson, S. M., Saygin, A. P., Sereno, M. I., & Iacoboni, M. (2004). Listening to speech activates motor areas involved in speech production. Nature Neuroscience, 7(7), 701--702.

Witteman, J., van Heuven, V. J., & Schiller, N. O. (2012). Hearing feelings: a quantitative meta-analysis on the neuroimaging literature of emotional prosody perception. Neuropsychologia, 50(12), 2752--2763.

Woolgar, A., & Zopf, R. (2017). Multisensory coding in the multiple-demand regions: Vibrotactile task information is coded in frontoparietal cortex. Journal of Neurophysiology, 118(2), 703--716. 10.1152/jn.00559.2016

Wu, Y., Uluç, I., Schmidt, T. T., Tertel, K., Kirilina, E., & Blankenburg, F. (2018). Overlapping frontoparietal networks for tactile and visual parametric working memory representations. NeuroImage, 166, 325--334.

Zaki, J., Davis, J. I., & Ochsner, K. N. (2012). Overlapping activity in anterior insula during interoception and emotional experience. NeuroImage, 62(1), 493--499.

Zora, H., Rudner, M., & Magnusson, A. K. M. (2020). Concurrent affective and linguistic prosody with the same emotional valence elicits a late positive ERP response. European Journal of Neuroscience, 51(11), 2236--2249.

