## Supplementary material for "Neural dynamics of induced throat vibrations during vocal emotion recognition"

**Contents table**

### Supplementary EEG methods

#### I. Event-related topographical clustering analysis

For each channel, the grand-average ERP for each condition was averaged into forty 25ms bins from voice onset to 1s post-stimulus onset. To increase the specificity of the topographical clustering algorithm to relevant temporal regions of interest, a baseline threshold was defined as ± 3 standard deviations of the grand average ERP for all conditions in the 1s baseline period. A database was created with, as lines the 40 timebins * 15 conditions (3 Vibration * 5 Morphing), and as columns the 64 EEG channels for a final matrix size of 600 lines * 64 columns = 38’400 cells. If the average power in the bin exceeded the baseline threshold, the average power was reported in the table and if the average power in the bin was within the baseline confidence interval, a 0 was reported in the database instead. Clustering used 25-ms bins; the statistical analyses used 50-ms bins (Methods 2.4.3).

Dimension reduction and topographical clustering was performed using Uniform Manifold Approximation and Projection (UMAP) for Dimension Reduction and Hierarchical Density-Based Spatial Clustering of Applications with Noise (HDBSCAN) for clustering both available in python (McInnes et al., 2017). Previous publications highlighted these methods as the most efficient for noisy high-dimensional data (Allaoui et al., 2020; McInnes et al., 2018; Weijler et al., 2022; Yang et al., 2021). UMAP algorithm was applied to the data with N_neighbors_=40 parameters. The number of components was chosen using a measure of specificity—the average proportion of components significantly correlated (Pearson’s test, *p<*.05) with each parameter, and inclusivity—the total proportion of parameters significantly correlated to at least one component. Using this method, the optimal number of components (highest inclusivity *and* specificity) for this dataset was determined to be 6.

The UMAP components were entered into a HDBSCAN algorithm with Cityblock metric. The clustering yielded four topographical clusters (C1 posterior, 21 electrodes; C2 centro-temporal, 18; C3 anterior-lateral frontal, 10; C4 frontocentral, 15) and no unassigned electrode (Supplementary Figure 2).

#### II. ERP statistical analysis

The models are described in Methods 2.4.3. In brms syntax, each full model was Amplitude ~ Vibration * Morphing + ParticipantGender + SpeakerGender + (1 | Participant) + (1 | Electrode), with a Gaussian likelihood and sum-coded factors; the reduced model omitted the Vibration × Morphing term. Bayes factors were obtained by bridge sampling. Evidence was labelled + (3.2 < K ≤ 10), ++ (10 < K ≤ 100) and +++ (K > 100), following the Jeffreys scale; the same cut-offs inverted label evidence for the model without the interaction (Supplementary Table 4). Convergence was checked for the full models (R-hat ≤ 1.012, bulk ESS ≥ 610, tail ESS ≥ 1016). Because the models include no trial-level random intercept, electrodes of the same trial are treated as independent; Bayes factors may therefore overstate the evidence and are best read as ordinal. Posterior contrasts (emmeans, reference = no vibration) were computed only in timebins with K > 3.2 and are reported as posterior medians with 95% HPD intervals (Supplementary Tables 5 and 6). Sensitivity to the prior was not assessed.

#### III. Source reconstruction and analysis

Source reconstruction used MNE-Python on the preprocessed, epoched data (Methods 2.4.2). The forward solution used the fsaverage template with a three-layer boundary-element model. An LCMV beamformer with unit-noise-gain weight normalisation was computed, using 0 to 1 s as the signal window and −1 to 0 s as the noise window, and applied to the epochs. Activity was extracted in the 360 labels of the HCP-MMP1 atlas (mean-flip; Glasser et al., 2016) and averaged, for each trial, within the eight 50-ms windows with a supported interaction in C4 (centres at 75, 325, 475, 525, 625, 775, 825 and 925 ms). We used a sequential analysis approach to detect significant vibration effects for each source label in each time-window with a supported interaction in C4.

For each label and window, amplitude was modelled with a Gaussian mixed model (glmmTMB) RATE ~ Morphing * Vibration + (1 | Participant), after excluding values beyond three interquartile ranges.

If, in a parcel and time window, Morphing × Vibration interaction reached significance, joint omnibus test was performed testing the effect of vibration for each morphing level (Benjamini–Hochberg method FDR correction for each omnibus test, 173 significant ROIs/Timewindows X 5 morphing levels = 865 tests). If joint-omnibus tests reached significance, cContrasts anger − none and fear − none within each morphing level were obtained with emmeans (no emmeans adjustment) and expressed as z ratios. The p-values of all these contrasts (360 labels × 8 windows × 5 morphing levels × 2 vibrations = 28,800 tests) were adjusted together with the Benjamini-Hochberg procedure; omnibus tests and anger-versus-fear contrasts were not part of this family. The 55 contrasts with pFDR<.05 are listed in Supplementary Table 9. Windows were chosen from the sensor-level results, so these tests localize the effects and do not confirm them independently.

### Supplementary EEG results

#### I. Other clusters

Supplementary Table 4 gives the Bayes factor for the Vibration × Morphing interaction in each cluster and timebin. Evidence for the interaction was found in one timebin in C3 (75 ms, K = 5.4), in none in C2, and in three in C1 (75, 825 and 925 ms; K > 100 in each, the same three timebins as in C4). Supplementary Table 6 gives the contrasts of each vibration against no vibration in those timebins: the 95% HPD interval excluded zero in 5 of 10 contrasts in C3 and in 24 of 30 in C1 (absolute differences 0.11-0.57 µV). C2 was not analysed further.

#### II. Cluster C4

Supplementary Table 5 lists the 80 contrasts (2 vibrations × 5 morphing levels × 8 timebins) summarised in Section 3.2.1. Fifty-four excluded zero, 29 for fear and 25 for anger vibration. The number of credible contrasts per timebin was 10 of 10 at 75 ms, 6 at 325 ms, 8 at 475 ms, 4 at 525 ms, 8 at 625 ms, 6 at 775 ms, 6 at 825 ms and 6 at 925 ms; per morphing level it was 13 (10% fear), 10 (40%), 10 (50%), 10 (60%) and 11 (90%). Contrasts between anger and fear vibration are not reported.

**Supplementary Tables**

**Supplementary Table 1**
Results of the analysis of the acoustic parameters of the voice stimuli as a function of Morphing with Bonferroni corrected pvalues.

| **Variables** | ***F*(4, 45)** | ***P*** | ***Corrected P*** | ***R^2^_marginal_*** | ***R^2^_conditional_*** |
| --- | --- | --- | --- | --- | --- |
| Amplitude | 1.04 | .40 | >.99 | .08 | .08 |
| Loudness | .25 | .91 | >.99 | .01 | .38 |
| Pitch | 2.44 | .06 | .48 | .06 | .72 |
| Roughness | .24 | .91 | >.99 | .02 | .02 |
| Spectral Centroid | 1.52 | .21 | >.99 | .05 | .60 |
| Spectral Slope | .45 | .77 | >.99 | .03 | .11 |
| Duration | 1.83 | .14 | >.99 | .13 | .13 |
| F1 frequency | .09 | .99 | >.99 | .00 | .81 |

LMMs were conducted on the fifty stimuli for each acoustic parameter with the factor Morphing (FA_10/90_, FA_40/60_, FA_50/50_, FA_60/40_, and FA_90/10_) and the speaker gender and identity as random effects.

**Supplementary Table 2**
Results of the analysis of the acoustic parameters of the vibration stimuli as a function of Vibration with Bonferroni corrected p-values.

| **Variables** | ***F*(1, 18)** | ***P*** | ***Corrected P*** | ***R^2^_marginal_*** | ***R^2^_conditional_*** |
| --- | --- | --- | --- | --- | --- |
| Amplitude | 2.10 | .16 | >.99 | .10 | .10 |
| Loudness | .26 | .62 | >.99 | .01 | .32 |
| Pitch | 3.89 | .07 | .52 | .07 | .66 |
| Roughness | 2.02 | .17 | >.99 | .10 | .10 |
| Spectral Centroid | 6.49 | <.05 | .17 | .17 | .51 |
| Spectral Slope | .02 | .88 | >.99 | .00 | .05 |
| Duration | 7.49 | <.05 | .11 | .28 | .28 |
| F1 frequency | .58 | .46 | >.99 | .01 | .79 |

LMMs were conducted on the twenty stimuli for each acoustic parameter with the factor Vibration (100% anger stimuli and 100% fear stimuli) and the actor’s gender (female vs male) as a random effect.

**Supplementary Table 3**

Results of the analysis of the reaction times of the participants and complementary effects for response data.

| **Effect (RT)** | **χ²** | **df** | ***p*** |
| --- | --- | --- | --- |
| Vibration | 3.70 | 2 | .16 |
| Morphing | .69 | 4 | .41 |
| Vibration * Morphing | 3.84 | 8 | .15 |
| **Effect (Response)** |  |  |  |
| Vibration | 8.27 | 2 | .016 |
| Morphing | 17.76 | 4 | .001 |
| Vibration * Morphing | 23.96 | 8 | .002 |
| Participant gender | 25.56 | 1 | <.0001 |
| Speaker gender | 15.09 | 1 | <.001 |

*Note:* χ², Chi squared; df, degrees of freedom; p, *p*-value.

**Supplementary Table 4**

Bayes factors for the Vibration * Morphing interaction, by electrode cluster and 50-ms timebin.

| **Timebin (ms)** | **C1** | | **C2** | | **C3** | | **C4** | |
| --- | --- | --- | --- | --- | --- | --- | --- | --- |
|  | **K** | **Evidence** | **K** | **Evidence** | **K** | **Evidence** | **K** | **Evidence** |
| 25 | 3.52 × 10^−15^ | −−− | 1.60 × 10^−16^ | −−− | 7.14 × 10^−14^ | −−− | 2.29 × 10^−14^ | −−− |
| 75 | **9.75 × 10^6^** | **+++** | 1.20 × 10^−17^ | −−− | **5.4** | **+** | **5.10 × 10^3^** | **+++** |
| 125 | 1.49 × 10^−10^ | −−− | 1.41 × 10^−20^ | −−− | 3.08 × 10^−16^ | −−− | 1.99 × 10^−8^ | −−− |
| 175 | 6.22 × 10^−13^ | −−− | 5.96 × 10^−18^ | −−− | 2.67 × 10^−7^ | −−− | 5.85 × 10^−15^ | −−− |
| 225 | 2.22 × 10^−16^ | −−− | 8.30 × 10^−20^ | −−− | 1.67 × 10^−16^ | −−− | 2.96 × 10^−17^ | −−− |
| 275 | 7.82 × 10^−8^ | −−− | 6.11 × 10^−18^ | −−− | 2.64 × 10^−13^ | −−− | 2.82 × 10^−7^ | −−− |
| 325 | 1.25 × 10^−3^ | −−− | 3.98 × 10^−20^ | −−− | 1.87 × 10^−15^ | −−− | **6.12 × 10^5^** | **+++** |
| 375 | 1.09 × 10^−6^ | −−− | 1.19 × 10^−18^ | −−− | 4.80 × 10^−10^ | −−− | 2.97 × 10^−3^ | −−− |
| 425 | 9.78 × 10^−18^ | −−− | 1.14 × 10^−18^ | −−− | 4.16 × 10^−16^ | −−− | 1.21 × 10^−14^ | −−− |
| 475 | 1.06 | ns | 5.71 × 10^−13^ | −−− | 2.99 × 10^−10^ | −−− | **1.06 × 10^12^** | **+++** |
| 525 | 7.81 × 10^−10^ | −−− | 4.71 × 10^−18^ | −−− | 1.90 × 10^−16^ | −−− | **10.9** | **++** |
| 575 | 1.53 × 10^−11^ | −−− | 3.94 × 10^−18^ | −−− | 0.26 | − | 9.42 × 10^−10^ | −−− |
| 625 | 7.59 × 10^−4^ | −−− | 5.14 × 10^−19^ | −−− | 2.56 × 10^−7^ | −−− | **170** | **+++** |
| 675 | 6.44 × 10^−14^ | −−− | 3.32 × 10^−20^ | −−− | 1.19 × 10^−7^ | −−− | 2.75 × 10^−17^ | −−− |
| 725 | 2.52 × 10^−17^ | −−− | 1.28 × 10^−20^ | −−− | 4.73 × 10^−14^ | −−− | 3.69 × 10^−15^ | −−− |
| 775 | 1.65 × 10^−3^ | −−− | 5.36 × 10^−20^ | −−− | 1.19 × 10^−5^ | −−− | **6.41 × 10^8^** | **+++** |
| 825 | **1.38 × 10^29^** | **+++** | 1.92 × 10^−21^ | −−− | 2.52 | ns | **1.09 × 10^27^** | **+++** |
| 875 | 5.22 × 10^−5^ | −−− | 2.78 × 10^−20^ | −−− | 3.22 × 10^−9^ | −−− | 9.24 × 10^−12^ | −−− |
| 925 | **4.25 × 10^7^** | **+++** | 4.79 × 10^−18^ | −−− | 2.79 × 10^−5^ | −−− | **1.66 × 10^5^** | **+++** |
| 975 | 3.19 × 10^−13^ | −−− | 3.44 × 10^−20^ | −−− | 3.76 × 10^−10^ | −−− | 2.57 × 10^−12^ | −−− |

**Note.** K is the Bayes factor comparing the full model (Vibration × Morphing interaction) with the model without the interaction, estimated by bridge sampling; one Bayesian Gaussian mixed model was fitted per cluster and timebin. Timebins are labelled by their centre (e.g., 75 = 50–100 ms after voice onset). C1–C4 are the electrode clusters shown in Supplementary Figure 2 (C1: 21, C2: 18, C3: 10, C4: 15 electrodes). Evidence for the interaction (Jeffreys scale): + substantial (3.2 < K ≤ 10), ++ strong (10 < K ≤ 100), +++ decisive (K > 100). ns: inconclusive (0.31 < K ≤ 3.2). Evidence for the model without the interaction: − (0.1 ≤ K < 0.31), −− (0.01 ≤ K < 0.1), −−− (K < 0.01). Shaded bold cells: K > 3.2. Number of timebins with K > 3.2: C1: 3; C2: 0; C3: 1; C4: 8.

**Supplementary Table 5**

Cluster C4: effect of anger and fear vibration relative to no vibration, per morphing level, in the timebins supported by the Bayes factors.

| **Timebin (ms)** | **Morphing level** | **Contrast** | **Estimate (µV)** | **95% HPD** | **Interval excludes 0** |
| --- | --- | --- | --- | --- | --- |
| **75** | **10% fear (90% anger)** | **Anger − none** | **0.198** | **[0.090, 0.308]** | **Yes** |
| **75** | **10% fear (90% anger)** | **Fear − none** | **−0.330** | **[−0.437, −0.223]** | **Yes** |
| **75** | **40% fear (60% anger)** | **Anger − none** | **0.374** | **[0.267, 0.482]** | **Yes** |
| **75** | **40% fear (60% anger)** | **Fear − none** | **0.299** | **[0.200, 0.407]** | **Yes** |
| **75** | **50% fear (50% anger)** | **Anger − none** | **0.113** | **[0.012, 0.222]** | **Yes** |
| **75** | **50% fear (50% anger)** | **Fear − none** | **0.131** | **[0.030, 0.240]** | **Yes** |
| **75** | **60% fear (40% anger)** | **Anger − none** | **0.230** | **[0.119, 0.330]** | **Yes** |
| **75** | **60% fear (40% anger)** | **Fear − none** | **0.208** | **[0.102, 0.311]** | **Yes** |
| **75** | **90% fear (10% anger)** | **Anger − none** | **0.107** | **[0.001, 0.216]** | **Yes** |
| **75** | **90% fear (10% anger)** | **Fear − none** | **0.150** | **[0.039, 0.254]** | **Yes** |
| 325 | 10% fear (90% anger) | Anger − none | −0.062 | [−0.174, 0.049] | No |
| **325** | **10% fear (90% anger)** | **Fear − none** | **−0.208** | **[−0.316, −0.095]** | **Yes** |
| **325** | **40% fear (60% anger)** | **Anger − none** | **−0.179** | **[−0.285, −0.073]** | **Yes** |
| **325** | **40% fear (60% anger)** | **Fear − none** | **−0.186** | **[−0.299, −0.086]** | **Yes** |
| 325 | 50% fear (50% anger) | Anger − none | −0.078 | [−0.183, 0.027] | No |
| **325** | **50% fear (50% anger)** | **Fear − none** | **−0.209** | **[−0.318, −0.106]** | **Yes** |
| 325 | 60% fear (40% anger) | Anger − none | −0.041 | [−0.144, 0.070] | No |
| 325 | 60% fear (40% anger) | Fear − none | 0.039 | [−0.072, 0.140] | No |
| **325** | **90% fear (10% anger)** | **Anger − none** | **0.249** | **[0.141, 0.363]** | **Yes** |
| **325** | **90% fear (10% anger)** | **Fear − none** | **0.496** | **[0.387, 0.603]** | **Yes** |
| **475** | **10% fear (90% anger)** | **Anger − none** | **−0.257** | **[−0.364, −0.152]** | **Yes** |
| **475** | **10% fear (90% anger)** | **Fear − none** | **−0.152** | **[−0.256, −0.044]** | **Yes** |
| **475** | **40% fear (60% anger)** | **Anger − none** | **0.109** | **[0.001, 0.213]** | **Yes** |
| **475** | **40% fear (60% anger)** | **Fear − none** | **−0.359** | **[−0.460, −0.248]** | **Yes** |
| **475** | **50% fear (50% anger)** | **Anger − none** | **−0.382** | **[−0.481, −0.269]** | **Yes** |
| **475** | **50% fear (50% anger)** | **Fear − none** | **−0.360** | **[−0.465, −0.254]** | **Yes** |
| **475** | **60% fear (40% anger)** | **Anger − none** | **0.147** | **[0.044, 0.254]** | **Yes** |
| 475 | 60% fear (40% anger) | Fear − none | 0.011 | [−0.089, 0.119] | No |
| 475 | 90% fear (10% anger) | Anger − none | −0.097 | [−0.200, 0.014] | No |
| **475** | **90% fear (10% anger)** | **Fear − none** | **0.112** | **[0.008, 0.222]** | **Yes** |
| **525** | **10% fear (90% anger)** | **Anger − none** | **−0.359** | **[−0.472, −0.259]** | **Yes** |
| **525** | **10% fear (90% anger)** | **Fear − none** | **−0.325** | **[−0.432, −0.218]** | **Yes** |
| 525 | 40% fear (60% anger) | Anger − none | 0.050 | [−0.051, 0.157] | No |
| 525 | 40% fear (60% anger) | Fear − none | −0.087 | [−0.189, 0.015] | No |
| 525 | 50% fear (50% anger) | Anger − none | 0.016 | [−0.092, 0.117] | No |
| 525 | 50% fear (50% anger) | Fear − none | −0.028 | [−0.127, 0.084] | No |
| 525 | 60% fear (40% anger) | Anger − none | 0.008 | [−0.096, 0.109] | No |
| **525** | **60% fear (40% anger)** | **Fear − none** | **−0.230** | **[−0.327, −0.113]** | **Yes** |
| **525** | **90% fear (10% anger)** | **Anger − none** | **−0.388** | **[−0.487, −0.281]** | **Yes** |
| 525 | 90% fear (10% anger) | Fear − none | −0.065 | [−0.171, 0.037] | No |
| **625** | **10% fear (90% anger)** | **Anger − none** | **−0.361** | **[−0.466, −0.258]** | **Yes** |
| **625** | **10% fear (90% anger)** | **Fear − none** | **−0.213** | **[−0.319, −0.107]** | **Yes** |
| 625 | 40% fear (60% anger) | Anger − none | −0.051 | [−0.155, 0.054] | No |
| **625** | **40% fear (60% anger)** | **Fear − none** | **−0.258** | **[−0.362, −0.148]** | **Yes** |
| 625 | 50% fear (50% anger) | Anger − none | −0.095 | [−0.200, 0.010] | No |
| **625** | **50% fear (50% anger)** | **Fear − none** | **−0.220** | **[−0.319, −0.114]** | **Yes** |
| **625** | **60% fear (40% anger)** | **Anger − none** | **0.155** | **[0.043, 0.254]** | **Yes** |
| **625** | **60% fear (40% anger)** | **Fear − none** | **0.202** | **[0.100, 0.304]** | **Yes** |
| **625** | **90% fear (10% anger)** | **Anger − none** | **−0.355** | **[−0.462, −0.247]** | **Yes** |
| **625** | **90% fear (10% anger)** | **Fear − none** | **−0.149** | **[−0.250, −0.039]** | **Yes** |
| **775** | **10% fear (90% anger)** | **Anger − none** | **0.238** | **[0.128, 0.342]** | **Yes** |
| **775** | **10% fear (90% anger)** | **Fear − none** | **0.253** | **[0.139, 0.357]** | **Yes** |
| **775** | **40% fear (60% anger)** | **Anger − none** | **−0.298** | **[−0.399, −0.187]** | **Yes** |
| 775 | 40% fear (60% anger) | Fear − none | 0.060 | [−0.045, 0.166] | No |
| 775 | 50% fear (50% anger) | Anger − none | −0.089 | [−0.200, 0.010] | No |
| **775** | **50% fear (50% anger)** | **Fear − none** | **0.196** | **[0.093, 0.303]** | **Yes** |
| 775 | 60% fear (40% anger) | Anger − none | 0.080 | [−0.022, 0.190] | No |
| 775 | 60% fear (40% anger) | Fear − none | −0.056 | [−0.157, 0.052] | No |
| **775** | **90% fear (10% anger)** | **Anger − none** | **−0.191** | **[−0.303, −0.088]** | **Yes** |
| **775** | **90% fear (10% anger)** | **Fear − none** | **−0.325** | **[−0.433, −0.220]** | **Yes** |
| 825 | 10% fear (90% anger) | Anger − none | 0.072 | [−0.036, 0.179] | No |
| 825 | 10% fear (90% anger) | Fear − none | 0.034 | [−0.069, 0.142] | No |
| **825** | **40% fear (60% anger)** | **Anger − none** | **−0.334** | **[−0.439, −0.229]** | **Yes** |
| 825 | 40% fear (60% anger) | Fear − none | 0.047 | [−0.060, 0.151] | No |
| **825** | **50% fear (50% anger)** | **Anger − none** | **0.459** | **[0.352, 0.561]** | **Yes** |
| **825** | **50% fear (50% anger)** | **Fear − none** | **0.330** | **[0.223, 0.435]** | **Yes** |
| **825** | **60% fear (40% anger)** | **Anger − none** | **−0.411** | **[−0.517, −0.297]** | **Yes** |
| **825** | **60% fear (40% anger)** | **Fear − none** | **−0.308** | **[−0.414, −0.201]** | **Yes** |
| **825** | **90% fear (10% anger)** | **Anger − none** | **0.196** | **[0.087, 0.308]** | **Yes** |
| 825 | 90% fear (10% anger) | Fear − none | −0.043 | [−0.150, 0.066] | No |
| **925** | **10% fear (90% anger)** | **Anger − none** | **−0.184** | **[−0.286, −0.079]** | **Yes** |
| **925** | **10% fear (90% anger)** | **Fear − none** | **−0.225** | **[−0.325, −0.118]** | **Yes** |
| 925 | 40% fear (60% anger) | Anger − none | 0.078 | [−0.024, 0.184] | No |
| **925** | **40% fear (60% anger)** | **Fear − none** | **0.241** | **[0.137, 0.342]** | **Yes** |
| 925 | 50% fear (50% anger) | Anger − none | 0.026 | [−0.078, 0.129] | No |
| **925** | **50% fear (50% anger)** | **Fear − none** | **0.331** | **[0.220, 0.430]** | **Yes** |
| **925** | **60% fear (40% anger)** | **Anger − none** | **−0.314** | **[−0.416, −0.210]** | **Yes** |
| **925** | **60% fear (40% anger)** | **Fear − none** | **−0.302** | **[−0.402, −0.196]** | **Yes** |
| 925 | 90% fear (10% anger) | Anger − none | 0.064 | [−0.039, 0.173] | No |
| 925 | 90% fear (10% anger) | Fear − none | 0.090 | [−0.014, 0.196] | No |

**Note.** Estimates are posterior medians of the difference in mean amplitude (µV) between each vibration condition and the no-vibration condition (positive: more positive amplitude with vibration), obtained with emmeans (treatment-versus-control, reference = no vibration) from the cluster C4 Bayesian model of each timebin. Only the eight timebins with K > 3.2 in C4 are shown (centres at 75, 325, 475, 525, 625, 775, 825, 925 ms; Supplementary Table 4). HPD: highest posterior density interval. Bold rows: the 95% HPD interval excludes zero (54 of 80 contrasts; absolute differences 0.107–0.496 µV). Morphing levels are expressed as percentage of fear (and anger) in the voice. Anger versus fear vibration contrasts are not reported.

**Supplementary Table 6**

Clusters C1 and C3: effect of anger and fear vibration relative to no vibration, per morphing level, in the timebins supported by the Bayes factors.

| **Cluster** | **Timebin (ms)** | **Morphing level** | **Contrast** | **Estimate (µV)** | **95% HPD** | **Interval excludes 0** |
| --- | --- | --- | --- | --- | --- | --- |
| **C3** | **75** | **10% fear (90% anger)** | **Anger − none** | **0.175** | **[0.050, 0.311]** | **Yes** |
| **C3** | **75** | **10% fear (90% anger)** | **Fear − none** | **−0.218** | **[−0.354, −0.090]** | **Yes** |
| **C3** | **75** | **40% fear (60% anger)** | **Anger − none** | **0.572** | **[0.442, 0.708]** | **Yes** |
| **C3** | **75** | **40% fear (60% anger)** | **Fear − none** | **0.485** | **[0.361, 0.613]** | **Yes** |
| C3 | 75 | 50% fear (50% anger) | Anger − none | 0.106 | [−0.025, 0.234] | No |
| **C3** | **75** | **50% fear (50% anger)** | **Fear − none** | **0.234** | **[0.105, 0.363]** | **Yes** |
| C3 | 75 | 60% fear (40% anger) | Anger − none | 0.014 | [−0.119, 0.145] | No |
| C3 | 75 | 60% fear (40% anger) | Fear − none | −0.020 | [−0.148, 0.109] | No |
| C3 | 75 | 90% fear (10% anger) | Anger − none | 0.043 | [−0.086, 0.173] | No |
| C3 | 75 | 90% fear (10% anger) | Fear − none | 0.101 | [−0.031, 0.234] | No |
| **C1** | **75** | **10% fear (90% anger)** | **Anger − none** | **−0.151** | **[−0.267, −0.036]** | **Yes** |
| **C1** | **75** | **10% fear (90% anger)** | **Fear − none** | **0.381** | **[0.265, 0.493]** | **Yes** |
| **C1** | **75** | **40% fear (60% anger)** | **Anger − none** | **−0.416** | **[−0.531, −0.298]** | **Yes** |
| **C1** | **75** | **40% fear (60% anger)** | **Fear − none** | **−0.396** | **[−0.508, −0.283]** | **Yes** |
| **C1** | **75** | **50% fear (50% anger)** | **Anger − none** | **−0.146** | **[−0.260, −0.031]** | **Yes** |
| **C1** | **75** | **50% fear (50% anger)** | **Fear − none** | **−0.157** | **[−0.266, −0.043]** | **Yes** |
| **C1** | **75** | **60% fear (40% anger)** | **Anger − none** | **−0.119** | **[−0.232, −0.011]** | **Yes** |
| **C1** | **75** | **60% fear (40% anger)** | **Fear − none** | **−0.151** | **[−0.256, −0.032]** | **Yes** |
| **C1** | **75** | **90% fear (10% anger)** | **Anger − none** | **−0.133** | **[−0.244, −0.016]** | **Yes** |
| **C1** | **75** | **90% fear (10% anger)** | **Fear − none** | **−0.201** | **[−0.310, −0.084]** | **Yes** |
| C1 | 825 | 10% fear (90% anger) | Anger − none | 0.033 | [−0.073, 0.148] | No |
| **C1** | **825** | **10% fear (90% anger)** | **Fear − none** | **0.112** | **[0.003, 0.222]** | **Yes** |
| **C1** | **825** | **40% fear (60% anger)** | **Anger − none** | **0.417** | **[0.298, 0.529]** | **Yes** |
| C1 | 825 | 40% fear (60% anger) | Fear − none | 0.088 | [−0.018, 0.206] | No |
| **C1** | **825** | **50% fear (50% anger)** | **Anger − none** | **−0.495** | **[−0.609, −0.383]** | **Yes** |
| **C1** | **825** | **50% fear (50% anger)** | **Fear − none** | **−0.316** | **[−0.424, −0.204]** | **Yes** |
| **C1** | **825** | **60% fear (40% anger)** | **Anger − none** | **0.410** | **[0.300, 0.520]** | **Yes** |
| **C1** | **825** | **60% fear (40% anger)** | **Fear − none** | **0.458** | **[0.346, 0.564]** | **Yes** |
| **C1** | **825** | **90% fear (10% anger)** | **Anger − none** | **−0.214** | **[−0.337, −0.104]** | **Yes** |
| C1 | 825 | 90% fear (10% anger) | Fear − none | −0.046 | [−0.155, 0.073] | No |
| C1 | 925 | 10% fear (90% anger) | Anger − none | 0.109 | [−0.008, 0.216] | No |
| **C1** | **925** | **10% fear (90% anger)** | **Fear − none** | **0.115** | **[0.012, 0.235]** | **Yes** |
| C1 | 925 | 40% fear (60% anger) | Anger − none | −0.040 | [−0.145, 0.073] | No |
| **C1** | **925** | **40% fear (60% anger)** | **Fear − none** | **−0.220** | **[−0.328, −0.108]** | **Yes** |
| C1 | 925 | 50% fear (50% anger) | Anger − none | −0.102 | [−0.208, 0.002] | No |
| **C1** | **925** | **50% fear (50% anger)** | **Fear − none** | **−0.221** | **[−0.331, −0.116]** | **Yes** |
| **C1** | **925** | **60% fear (40% anger)** | **Anger − none** | **0.385** | **[0.279, 0.500]** | **Yes** |
| **C1** | **925** | **60% fear (40% anger)** | **Fear − none** | **0.476** | **[0.367, 0.582]** | **Yes** |
| **C1** | **925** | **90% fear (10% anger)** | **Anger − none** | **−0.180** | **[−0.287, −0.063]** | **Yes** |
| **C1** | **925** | **90% fear (10% anger)** | **Fear − none** | **−0.169** | **[−0.279, −0.053]** | **Yes** |

**Note.** Same conventions as Supplementary Table 5. Only the timebins with K > 3.2 are shown: C3 at 75 ms, and C1 at 75, 825 and 925 ms (Supplementary Table 4). No timebin in C2 showed evidence for the interaction. Bold rows: the 95% HPD interval excludes zero (29 of 40 contrasts; absolute differences 0.112–0.572 µV). Anger versus fear vibration contrasts are not reported.

**Supplementary Table 7**

Average trials (± Standard deviation) per conditions/channels/participants remaining after artifacts rejection (EEG data)

| **Vibration/Emotion** | **FA_10/90_** | **FA_40/60_** | **FA_50/50_** | **FA_60/40_** | **FA_90/10_** |
| --- | --- | --- | --- | --- | --- |
| **Anger** | 39.83±5.08 | 41.25±6.69 | 42.17±5.55 | 41.17±5.15 | 40.46±5.14 |
| **None** | 40.25±7.45 | 40.63±5.91 | 41.33±6.69 | 42.25±7.25 | 39.58±4.68 |
| **Fear** | 40.42±4.23 | 41.46±7.11 | 41.92±6.05 | 41.21±6.69 | 40.42±5.75 |

Note: Voice stimuli morphed 90% anger – 10% fear (FA_10/90_), 60% anger – 40% fear (FA_40/60_), 50% fear (FA_50/50_), 40% anger – 60% fear (FA_60/40_), 10% anger – 90% fear (FA_90/10_).

**Supplementary Table 8**

Effect of anger and fear vibration relative to no vibration on the odds of a fear response, by morphing level.

| **Morphing level** | **Contrast** | **OR** | **SE** | **95% CI** | **z** | **p** |
| --- | --- | --- | --- | --- | --- | --- |
| 10% fear (90% anger) | Anger / No Vibration | 1.224 | 0.1430 | [0.945, 1.586] | 1.733 | .1513 |
| **10% fear (90% anger)** | **Fear / No Vibration** | **1.449** | **0.1600** | **[1.134, 1.852]** | **3.359** | **.0015** |
| 40% fear (60% anger) | Anger / No Vibration | 1.125 | 0.1150 | [0.896, 1.413] | 1.149 | .4129 |
| **40% fear (60% anger)** | **Fear / No Vibration** | **1.336** | **0.1300** | **[1.076, 1.658]** | **2.972** | **.0058** |
| **50% fear (50% anger)** | **Anger / No Vibration** | **0.764** | **0.0786** | **[0.608, 0.960]** | **−2.617** | **.0172** |
| 50% fear (50% anger) | Fear / No Vibration | 0.948 | 0.0929 | [0.763, 1.179] | −0.540 | .7987 |
| **60% fear (40% anger)** | **Anger / No Vibration** | **0.772** | **0.0818** | **[0.610, 0.977]** | **−2.443** | **.0280** |
| 60% fear (40% anger) | Fear / No Vibration | 1.026 | 0.1050 | [0.818, 1.288] | 0.252 | .9426 |
| 90% fear (10% anger) | Anger / No Vibration | 0.838 | 0.1030 | [0.637, 1.102] | −1.433 | .2648 |
| 90% fear (10% anger) | Fear / No Vibration | 0.941 | 0.1130 | [0.720, 1.229] | −0.505 | .8192 |

**Note.** OR: odds ratio of a ‘fear’ response, vibration relative to no vibration (OR > 1: more fear responses with vibration). SE: standard error of the OR. 95% CI: confidence interval of the OR, back-transformed from the log odds ratio scale. z: Wald z ratio (df = ∞). Estimates are from the interaction of the binomial mixed model (emmeans), averaged over participant gender and speaker gender. P-value and confidence-interval adjustment: Dunnett method for 2 tests (anger vibration vs no vibration; fear vibration vs no vibration), within each morphing level. Tests are performed on the log odds ratio scale. Bold rows: adjusted p < .05.

**Supplementary Table 9**

Source-level effects of anger and fear vibration relative to no vibration surviving false-discovery-rate correction.

| **Window (ms)** | **Morphing level** | **Contrast** | **Hem.** | **ROI** | **Region (HCP-MMP1)** | **z** | **pFDR** |
| --- | --- | --- | --- | --- | --- | --- | --- |
| 50–100 | 10% fear (90% anger) | Anger − none | R | H | Hippocampus | −3.57 | .002 |
| 50–100 | 10% fear (90% anger) | Anger − none | R | PHA1 | ParaHippocampal Area 1 | −3.57 | .002 |
| 50–100 | 10% fear (90% anger) | Anger − none | R | PHA2 | ParaHippocampal Area 2 | −3.55 | .002 |
| 50–100 | 10% fear (90% anger) | Anger − none | R | PHA3 | ParaHippocampal Area 3 | −3.30 | .003 |
| 50–100 | 10% fear (90% anger) | Anger − none | R | VMV1 | VentroMedial Visual Area 1 | −3.64 | .002 |
| 50–100 | 10% fear (90% anger) | Anger − none | R | VMV2 | VentroMedial Visual Area 2 | −3.68 | .001 |
| 50–100 | 10% fear (90% anger) | Fear − none | R | H | Hippocampus | −2.20 | .045 |
| 50–100 | 10% fear (90% anger) | Fear − none | R | PHA1 | ParaHippocampal Area 1 | −2.55 | .020 |
| 50–100 | 10% fear (90% anger) | Fear − none | R | PHA2 | ParaHippocampal Area 2 | −2.43 | .027 |
| 50–100 | 10% fear (90% anger) | Fear − none | R | PHA3 | ParaHippocampal Area 3 | −2.76 | .012 |
| 50–100 | 10% fear (90% anger) | Fear − none | R | VMV1 | VentroMedial Visual Area 1 | −2.33 | .034 |
| 50–100 | 10% fear (90% anger) | Fear − none | R | VMV2 | VentroMedial Visual Area 2 | −2.18 | .046 |
| 50–100 | 40% fear (60% anger) | Anger − none | L | 3a | Area 3a | −3.71 | .001 |
| 50–100 | 50% fear (50% anger) | Anger − none | R | PHT | Area PHT | 3.35 | .002 |
| 50–100 | 50% fear (50% anger) | Anger − none | R | TPOJ1 | Area TemporoParietoOccipital Junction 1 | 3.44 | .002 |
| 50–100 | 50% fear (50% anger) | Fear − none | R | PHA1 | ParaHippocampal Area 1 | −3.22 | .003 |
| 50–100 | 50% fear (50% anger) | Fear − none | R | ProS | ProStriate Area | −3.49 | .002 |
| 50–100 | 50% fear (50% anger) | Fear − none | R | VMV1 | VentroMedial Visual Area 1 | −3.35 | .002 |
| 50–100 | 60% fear (40% anger) | Anger − none | R | TPOJ2 | Area TemporoParietoOccipital Junction 2 | 3.61 | .002 |
| 50–100 | 60% fear (40% anger) | Fear − none | R | 6mp | Area 6mp (mesial posterior) | −3.47 | .002 |
| 50–100 | 60% fear (40% anger) | Fear − none | R | TPOJ2 | Area TemporoParietoOccipital Junction 2 | 2.75 | .012 |
| 50–100 | 90% fear (10% anger) | Fear − none | R | PHT | Area PHT | −2.96 | .007 |
| 50–100 | 90% fear (10% anger) | Fear − none | R | TE1p | Area TE1 Posterior | −2.82 | .010 |
| 50–100 | 90% fear (10% anger) | Fear − none | R | TPOJ2 | Area TemporoParietoOccipital Junction 2 | −3.04 | .006 |
| 300–350 | 10% fear (90% anger) | Anger − none | R | VMV3 | VentroMedial Visual Area 3 | −3.47 | .002 |
| 300–350 | 10% fear (90% anger) | Fear − none | R | VMV3 | VentroMedial Visual Area 3 | −2.73 | .013 |
| 300–350 | 10% fear (90% anger) | Fear − none | L | ProS | ProStriate Area | −2.89 | .008 |
| 300–350 | 50% fear (50% anger) | Fear − none | R | VMV1 | VentroMedial Visual Area 1 | −3.43 | .002 |
| 300–350 | 60% fear (40% anger) | Anger − none | R | VMV3 | VentroMedial Visual Area 3 | −3.47 | .002 |
| 450–500 | 60% fear (40% anger) | Fear − none | R | 13l | Area 13l | −2.69 | .014 |
| 450–500 | 60% fear (40% anger) | Fear − none | R | EC | EntoRhinal Cortex | −3.50 | .002 |
| 450–500 | 60% fear (40% anger) | Fear − none | R | FOP4 | Frontal Opercular Area 4 | −3.62 | .002 |
| 450–500 | 60% fear (40% anger) | Fear − none | R | MI | Middle Insular Area | −3.84 | .001 |
| 450–500 | 60% fear (40% anger) | Fear − none | R | PeEc | Perirhinal Ectorhinal Cortex | −3.95 | .001 |
| 450–500 | 60% fear (40% anger) | Fear − none | R | Pir | Piriform Cortex | −3.81 | .001 |
| 450–500 | 60% fear (40% anger) | Fear − none | R | TE2a | Area TE2 Anterior | −3.78 | .001 |
| 450–500 | 60% fear (40% anger) | Fear − none | R | TF | Area TF | −3.56 | .002 |
| 450–500 | 60% fear (40% anger) | Fear − none | R | TGd | Area TG dorsal | −4.39 | <.001 |
| 450–500 | 60% fear (40% anger) | Fear − none | R | TGv | Area TG Ventral | −4.36 | <.001 |
| 450–500 | 60% fear (40% anger) | Fear − none | R | pOFC | posterior OFC Complex | −3.41 | .002 |
| 750–800 | 40% fear (60% anger) | Fear − none | R | p24 | Area posterior 24 | 2.62 | .017 |
| 900–950 | 10% fear (90% anger) | Anger − none | R | LIPd | Area Lateral IntraParietal dorsal | 2.71 | .013 |
| 900–950 | 10% fear (90% anger) | Anger − none | R | MIP | Medial IntraParietal Area | 2.91 | .008 |
| 900–950 | 10% fear (90% anger) | Anger − none | R | VIP | Ventral IntraParietal Complex | 3.52 | .002 |
| 900–950 | 10% fear (90% anger) | Fear − none | R | LIPd | Area Lateral IntraParietal dorsal | 3.23 | .003 |
| 900–950 | 10% fear (90% anger) | Fear − none | R | MIP | Medial IntraParietal Area | 3.08 | .005 |
| 900–950 | 10% fear (90% anger) | Fear − none | R | VIP | Ventral IntraParietal Complex | 2.28 | .038 |
| 900–950 | 50% fear (50% anger) | Fear − none | R | 7Am | Area 7Am | 3.64 | .002 |
| 900–950 | 60% fear (40% anger) | Anger − none | R | 46 | Area 46 | 3.54 | .002 |
| 900–950 | 60% fear (40% anger) | Anger − none | R | 9-46d | Area 9-46 dorsal | 3.89 | .001 |
| 900–950 | 60% fear (40% anger) | Anger − none | R | 9p | Area 9 Posterior | 4.32 | <.001 |
| 900–950 | 60% fear (40% anger) | Anger − none | R | d32 | Area dorsal 32 | 4.04 | .001 |
| 900–950 | 60% fear (40% anger) | Anger − none | R | p24 | Area posterior 24 | 3.68 | .001 |
| 900–950 | 60% fear (40% anger) | Fear − none | R | 9p | Area 9 Posterior | 2.56 | .019 |
| 900–950 | 60% fear (40% anger) | Fear − none | R | d32 | Area dorsal 32 | 2.23 | .043 |

**Note.** Source-level effects of vibration relative to no vibration surviving false-discovery-rate correction (*p*FDR<.05): 55 contrasts (20 anger, 35 fear; 53 in the right hemisphere). z is the z ratio of the emmeans contrast from a Gaussian mixed model fitted for each HCP-MMP1 label and 50-ms window (positive: higher source amplitude with vibration than without; negative: lower). For fear vibration the sign is that of the displayed contrast in Figure 4 (fear − none). *p*FDR: Benjamini-Hochberg adjusted p-value, computed across all ROI × window × morphing level × vibration contrasts; values below .001 are shown as <.001. Tested windows were the eight 50-ms timebins with a supported interaction in C4 (50-100, 300-350, 450-500, 500-550, 600-650, 750-800, 800-850 and 900-950 ms). Effects survived correction in five of them (50-100, 300-350, 450-500, 750-800 and 900-950 ms) and in none of the others. Hem.: hemisphere (L left, R right). ROI: HCP-MMP1 label (Glasser et al., 2016). Windows were chosen on the basis of the sensor-level results, so these tests localise the effects and are not independent confirmations. ‘non’: no-vibration condition.

**Supplementary Figures**

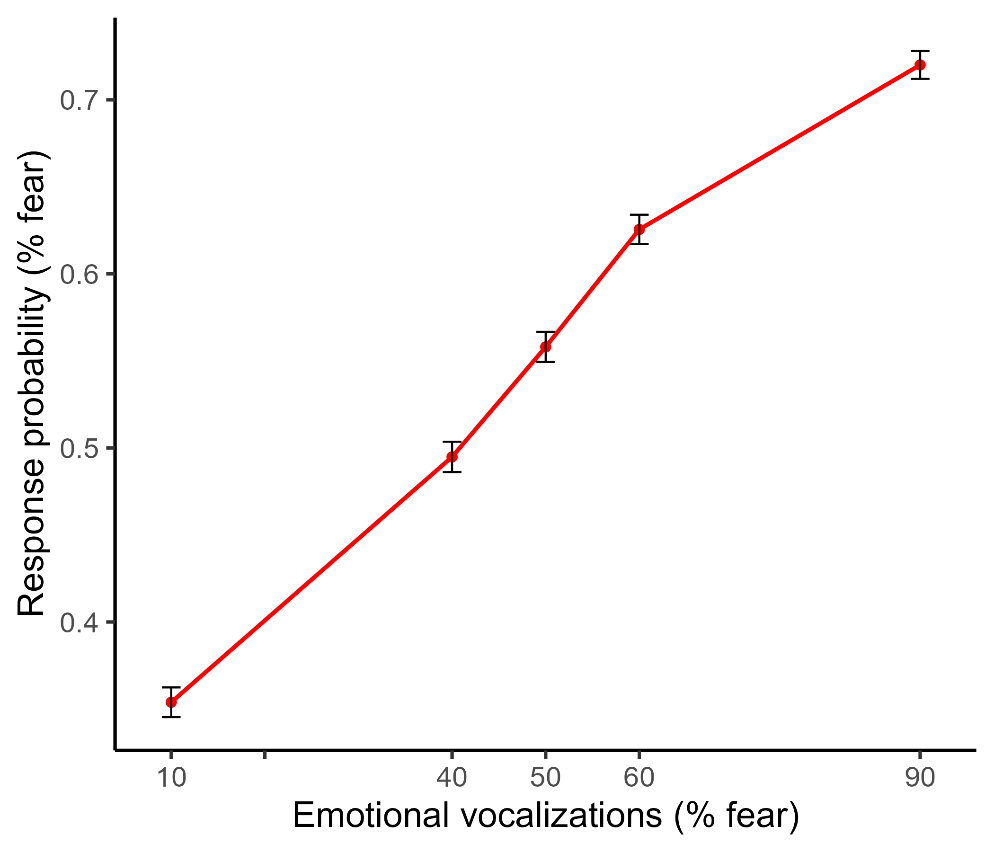

***Supplementary Figure 1***

**Main effect of Morphing on the probability of fear responses across vibration types.** On the X axis, each point represents a Morphing condition (i.e., FA_10/90_, FA_40/60_, FA_50/50_, FA_60/40_, FA_90/10_).

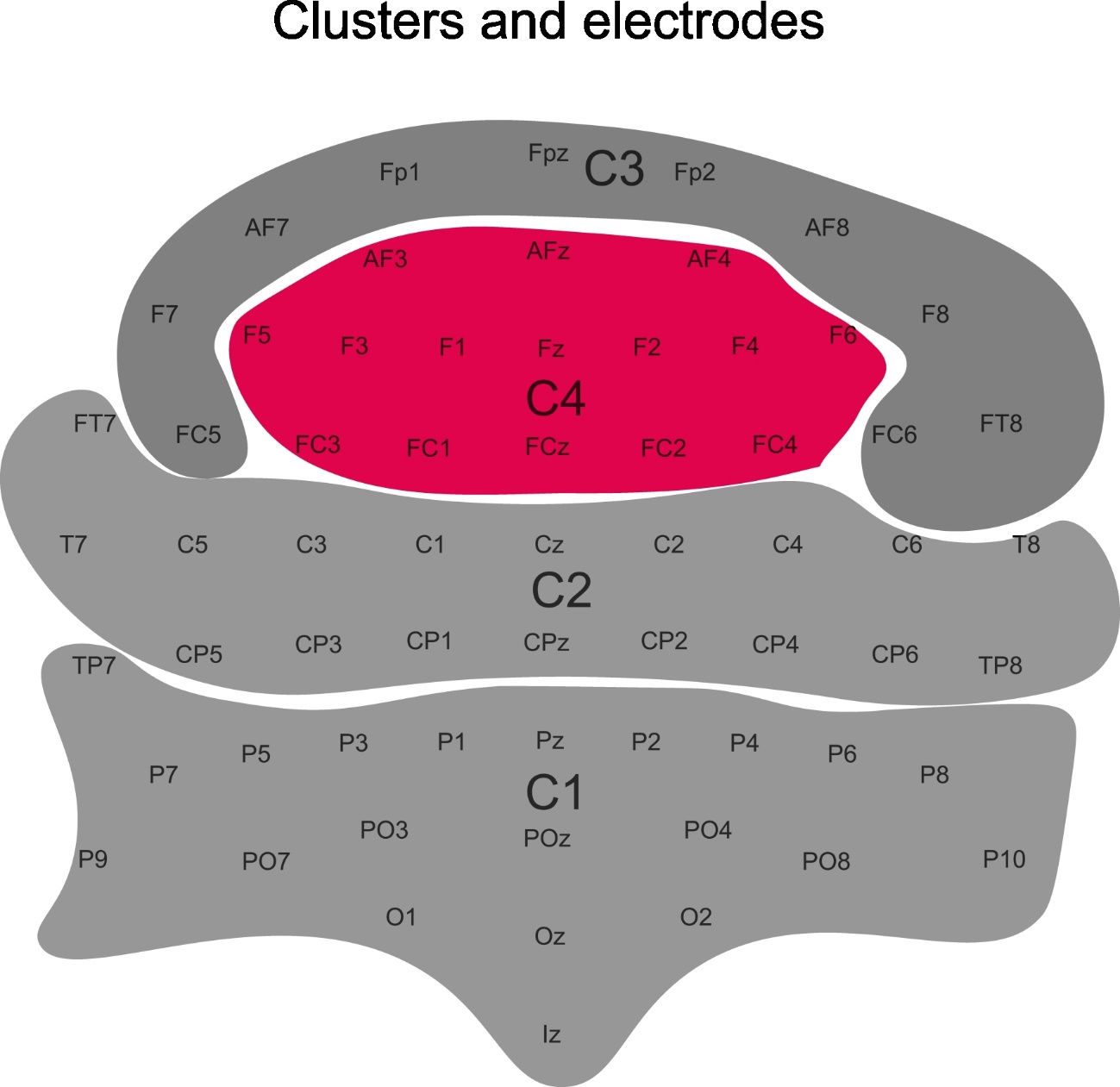

***Supplementary Figure 2***

**Topographical clusters of the 64 EEG electrodes.** Electrodes were grouped by UMAP dimension reduction and HDBSCAN clustering of the grand-average ERP pooled across all fifteen conditions (Supplementary EEG methods I), which yielded four clusters and no unassigned electrode: C1 (posterior, 21 electrodes), C2 (centro-temporal, 18), C3 (anterior-lateral frontal, 10) and C4 (frontocentral, 15, shown in red). Each cluster served as a region of interest in the ERP analyses (Methods 2.4.3). The cluster labels (C1–C4) are not electrode names: electrodes C1–C4 belong to cluster C2. The layout is schematic and not to scale; left–right differences in membership (e.g. FT7 in C2 and FT8 in C3) result from the data-driven clustering.

**References**

Allaoui, M., Kherfi, M. L., & Cheriet, A. (2020). Considerably improving clustering algorithms using UMAP dimensionality reduction technique: a comparative study. International conference on image and signal processing,

Glasser, M. F., Coalson, T. S., Robinson, E. C., Hacker, C. D., Harwell, J., & Yacoub, E. (2016). A multi-modal parcellation of human cerebral cortex. *Nature*, 1--11. https://doi.org/10.1038/nature18933

McInnes, L., Healy, J., & Astels, S. (2017). hdbscan: Hierarchical density based clustering. *Journal of Open Source Software*, *2*(11), 205.

McInnes, L., Healy, J., & Melville, J. (2018). UMAP: Uniform manifold approximation and projection for dimension reduction. In.

Weijler, L., Kowarsch, F., Wödlinger, M., Reiter, M., Maurer-Granofszky, M., Schumich, A., & Dworzak, M. N. (2022). UMAP based anomaly detection for minimal residual disease quantification within acute myeloid leukemia. *Cancers*, *14*(4), 898.

Yang, Y., Sun, H., Zhang, Y., Zhang, T., Gong, J., Wei, Y., & Yu, D. (2021). Dimensionality reduction by UMAP reinforces sample heterogeneity analysis in bulk transcriptomic data. *Cell Reports*, *36*(4).
